# Deep unfolded convolutional dictionary learning for motif discovery

**DOI:** 10.1101/2022.11.06.515322

**Authors:** Shane Chu, Gary Stormo

## Abstract

We present a principled representation learning approach based on convolutional dictionary learning (CDL) for motif discovery. We unroll an iterative algorithm that optimizes CDL as a forward pass in a neural network, resulting in a network that is fully interpretable, fast, and capable of finding motifs in large datasets. Simulated data show that our network is more sensitive and specific for discovering binding sites that exhibit complex binding patterns than popular motif discovery methods such as STREME and HOMER. Our network reveals statistically significant motifs and their diverse binding modes from the JASPAR database that are currently not reported.

## 1 Background and motivation

Finding conserved substrings in a set of unaligned DNA strings is a fundamental problem in computational biology. These conserved substrings, called *motifs*, arise from the convergence of evolutionary forces and are marked as molecular switches for regulatory mechanisms. Accurate characterization of motifs reveals binding specificity of proteins, which in turn provides features for downstream regulatory genomics tasks and helps understand how cells carry out gene expressions.

All models devised for motif discovery represent motifs in either a localist representation or a distributed representation [Hin84]. Most combinatorial and probabilistic formulations for motif discovery represent the motif in a localist representation – either as a consensus string, a product-multinomial distribution, or a position weight matrix (PWM).

While a localist representation is easy to interpret, the models that employ it to represent motifs tend to be overly simplistic, e.g., MEME [BE95] formulates motif discovery as a two-component mixture model, for which one component represents the motif, and the other component represents the genomic background. To overcome this over-simplification, e.g., so that MEME can find multiple motifs in the input strings, MEME performs sequential motif discovery: it “masks out” the substrings used by the discovered motifs before finding other motifs. Such an attempt is problematic for motif discovery as there may be more than one motif in the dataset, and motifs can exhibit binding modes such as *multi-meric binding* or *alternative structural conformations* [SG14]. Both STREME and HOMER still employ similar heuristics for motif discovery at the time of writing. The results section 3.3 shows how this heuristic can lead to incomplete motif discovery results.

The most popular model that employs distributed representation for motif discovery at the time of writing is the convolutional neural network (CNN). Although the model is expressive, it is challenging to interpret what represents the motif as the model itself is in a hierarchical distributed representation. Some attempts seek to interpret the first convolution layer of the CNN as motifs, e.g., [KP21; Nov+22], but provide no quantitative justifications for the choice of CNN architectures. Other attempts seek to find interpretations through model agnostic explanation methods such as SHAP [LL17]; however, SHAP is computationally expensive when the model contains many features.

To find an expressive, interpretable model that avoids the need for heuristics such as substring-masking, we ask the following question: what is a simple distributed representation on the input strings that we can use to infer the motifs? We find a solution to such a question to be convolutional dictionary learning (CDL) [GW18; Woh17; DI18; HHW15; BEL13]. The basic idea is we approximate each DNA string ***s*** as a sum of the convolution of feature vectors and sparse vectors, i.e.,

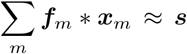

Conventionally, the feature vectors ***f***_*m*_’s are called the filters, and the sparse vectors ***x***’s are called the sparse code.

In this work, we present a motif discovery method based on CDL. The interpretation is straightforward: the CDL approximates each input sequence with a sparse linear combination of shift-invariant filters. This built-in interpretation can be efficiently used to infer the enriched patterns in the dataset, i.e., the motifs. Further, CDL allows us to *simultaneously* discover motifs in the input strings as opposed to sequential motif discovery. To scale CDL for large-scale motif discovery, we unfold the iterative procedure that optimizes CDL as a forward pass of a neural network [GL10; MLE21]. This results in a fully interpretable network that learns an approximate sparse representation in a timely manner. Our method discovers zero or more motifs for each sequence in the dataset. Simulated data suggests that our method is more capable of capturing multiple motifs in the dataset, especially when they share similar patterns. Additionally, our method reveals alternative modes of binding that are not reported in the JASPAR database [Cas+22].

## 2 Methodology

### 2.1 Notation

The term DNA strings referred to strings defined on the alphabet {A,C,G,T}. The map r(·) takes a vector and reshapes it into a matrix following the column-major order. The *i*-th component of a vector ***v*** is denoted as ***v***[*i*], and the *i*-th to the *j*-th consecutive components of ***v*** is denoted by ***v***[*i*:*j*]. Similarly, ***W*** [:, *i*] is the *i*-th column of the matrix ***W***. We write ***x*** ≥ **0** to indicate that all the components of the vector ***x*** are nonnegative. Brackets around an integer *K*, i.e., [*K*], is the set of integers {1, …, *K*}. The function | · | takes a finite set and returns its number of elements. The function ⌈·⌉ rounds up a real number. The map diag(·) returns a diagonal matrix with identical diagonal components from the input matrix. The norms ∥ · ∥_*F*_, ∥ · ∥_2_, and ∥ · ∥_1_ are the Frobenius norm, *ℓ*_2_ norm, and *ℓ*_1_ norm, respectively.

For the motif discovery problem, assume we have a set of one-hot encoded DNA strings ***s***_1_, …, ***s***_*N*_ each consisting of *L* nucleotides. Each string ***s***_*n*_ may contain zero or multiple motifs. We will first formulate convolutional dictionary learning (CDL) to obtain a sparse representation of the dataset. The sparse representation refers to a dictionary ***D*** which consists of filters ***d***_1_, …, ***d***_*M*_ and a sparse code ***X*** of (sparse) code vectors ***x***_11_, …, ***x***_*MN*_ (Figure 1 correspond to one DNA string, i.e., *N* = 1). Our goal with CDL is to obtain the sparse representation for which we could infer what and where the motifs are in the strings ***s***_1_, …, ***s***_*N*_.

**Figure 1:**
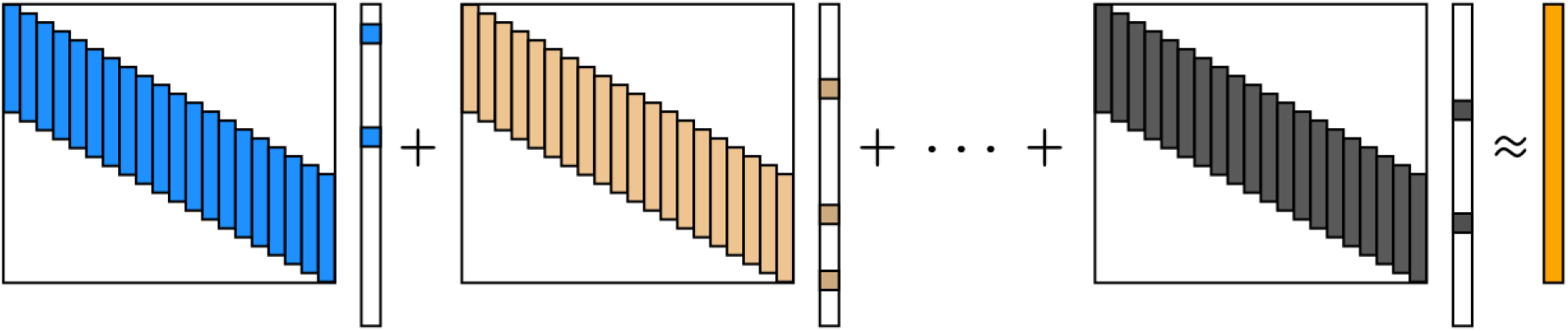
Approximate the signal (in orange) using a sum of convolutions. A convolution ***f*** * ***x*** is equivalent to ***F x***, a Toeplitz matrix ***F*** multiply a (sparse) vector ***x***.

**Figure 2:**
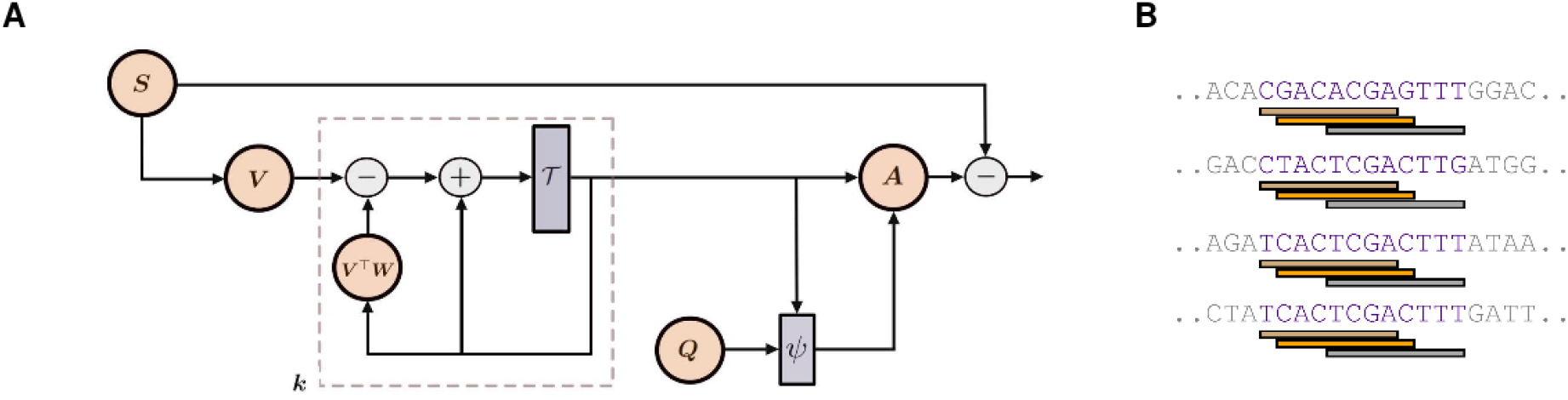
**A**. The computation graph of the unfolded CDL. Its forward pass corresponds to the alternating minimization of the CDL problem in 2. The dashed part corresponds to *k* ISTA iterations in CSC. The network then executes a mirror descent step with the distance function *ψ* for the dictionary update. **B**. Illustration of a triplet (equation 10): three filters, *m*_1_ in brown, *m*_2_ in orange, and *m*_3_ in grey that jointly represents a set of substrings (in purple) in a particular configuration.

### 2.2 Convolutional dictionary learning

We approximate each DNA sequence ***s*** as a sum of convolutions:

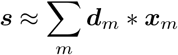

where ***d***_*m*_’s are the filters, and ***x***_*m*_’s are the code vectors. We consider the reshaped filters as position frequency matrices (PFM), i.e., r(***d***_*m*_) ∈ 𝒫_*ℓ*_, where

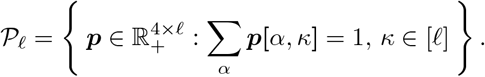

is the set of PFMs with *ℓ* columns, which avoids the scaling ambiguity between the filters and the sparse code. Now, with DNA strings ***s***_1_, …, ***s***_*N*_, we formulate the sparse representation discovery for DNA sequences as a convolutional dictionary learning (CDL) problem

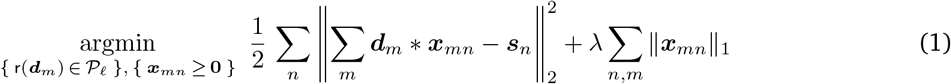

for which we can compactly write as

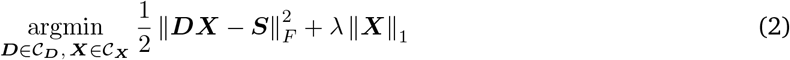

where matrices ***D, X, S*** are the concatenation of all the filters as Toeplitz matrices, all the code vectors, and all the input DNA sequences, respectively. The sets 𝒞_***D***_ and 𝒞_***X***_ encode the same constraints as in 1.^1^. The parameter *λ* controls the trade-off between the sparsity of ***X*** and the reconstruction loss.

### 2.3 Alternating minimization for convolutional dictionary learning

A common strategy to obtain the solution to problem 2 is via alternating minimization [GW18]. The alternating minimization proceeds by iteratively solving the two following convex sub-problems of 2:

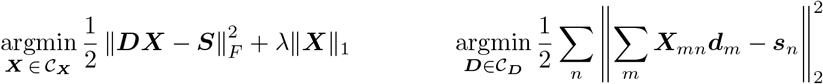

Here, ***X***_*mn*_ is the Toeplitz matrix that corresponds to ***x***_*mn*_. The problem on the left is convolution sparse coding (CSC), and the problem on the right is dictionary update (DU). It is well-known that CSC can be solved by the iterative shrinkage-thresholding algorithm (ISTA). Specifically, the sparse code ***X*** at the *t*-th iteration of ISTA is

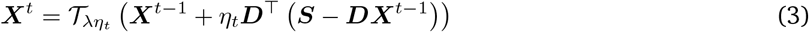

where *η*_*t*_ is the step-size at the *t*-th iteration, and 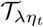 is the non-negative soft-thresholding operator parameterized by *λη*_*t*_. For DU, we consider mirror descent [NY83; BT03]. Let ***d***_*mκ*_ = ***d***_*m*_[4(*κ* − 1) + 1:4(*κ* − 1) + 4] be the *κ*-th non-overlapping four-consecutive components of ***d***_*m*_ and *ϕ*(***v***) = ∈_*i*_ ***v***_*i*_log***v***_*i*_ the negative entropy. We define the distance function associated with the Bregman divergence as

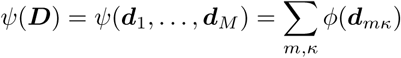

so an update step in mirror descent can be expressed as

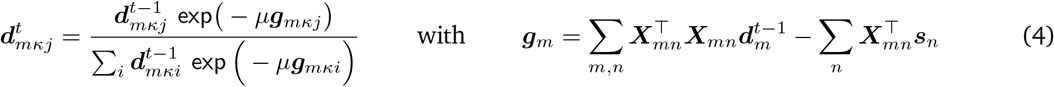

where ***g***_*m*_ is the gradient of ***d***_*m*_, *μ* is the step-size, and ***d***_*mκj*_ = ***d***_*mκ*_[*j*] is the *j*-th component of ***d***_*mκ*_. Informed by equations 3 and 4, we unfold this iterative procedure of alternating minimization into a neural network. This technique, called *deep unfolding*, obtains an approximate sparse representation and greatly speeds up the optimization [MLE21; GL10].

### 2.4 Approximate the sparse representation with deep unfolding

Our first *K* layers of the neural network consist of nonlinearities that correspond to the ISTA iterations in CSC:

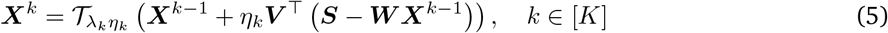

where the dictionary ***D*** in 3 is now “untied” and replaced with network weights ***V*** and ***W***. The untying of weights induces competition among the filters and their corresponding sparse codes, a phenomenon termed “mutual inhibition and explaining away” in [GL10]. The parameters *λ*_*k*_ and *η*_*k*_ now become learnable weights in the network. We take the sparse code from the *K*-th layer, ***X***^*K*^, and introduce the network weight ***Q*** that corresponds to filters ***q***_1_, …, ***q***_*M*_ to form a single update step to form a dictionary ***A***

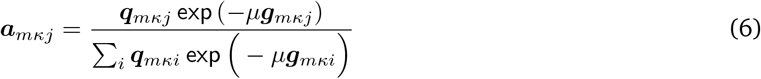

for which we keep the same notation convention as in 4, and the step-size *μ* is learnable. Altogether, the learnable parameter in the network is **Θ** = {***V***, ***W***, ***Q, η, λ***, *μ*} where ***η*** = {*η*_1_, …, *η*_*K*_} and ***λ*** = {*λ*_1_, …, *λ*_*K*_}. We formulate the loss of this network as

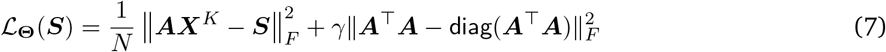

The term 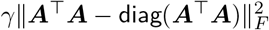 is the coherence regularizer that ensures the filters in ***A*** are diverse, and *γ* is a trade-off parameter of this regularizer and the reconstruction loss.

### 2.5 Motif discovery via sparse representations

Once we obtained the learned sparse representation from 7, we consider the set of sparse code components with high magnitude:

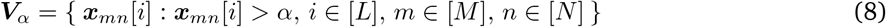

where *α* is a positive real number, e.g., selected by fixing a percentile of the magnitudes of all sparse-code components. Next, given three filters indexed by *m*_1_, *m*_2_, *m*_3_, integers △_1_, △_2_, △_3_, and a neighborhood parameter *δ*, we look for integers 1 ≤ *i*_1_ ≤ *i*_2_ ≤ *i*_3_ ≤ *L* and *n* ∈ [*N*] satisfying the following condition

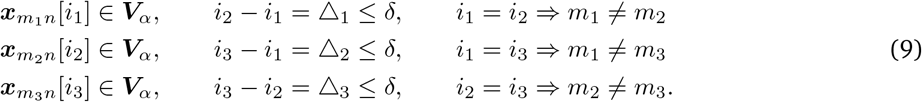

The set

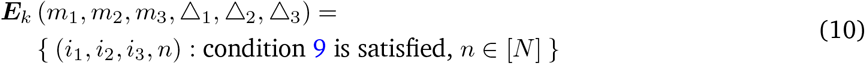

characterizes how three filters *m*_1_, *m*_2_, *m*_3_ jointly represent a set of DNA strings in a particular configuration. We will simply call such a set a *triplet* and its cardinality |***E***_*k*_ (*m*_1_, *m*_2_, *m*_3_, △_1_, △_2_, △_3_)| as *triplet count*.

The distribution of the triplet count is positively skewed. The highly enriched triplets on the right tend to represent the frequently occurring DNA sequence pattern, shown in figure 3. Each of these enriched triplets defines a multiple sequence alignment (MSA) and hence a PWM. Note that the enumeration of the triplet (equation 10), if done naively, results in time complexity O(*NL*^3^). However, when the valid neighborhoodparameter *δ* is small, e.g., *δ* ≤ 20, we can first locate the valid neighborhood of all the triplets with a binary search and then perform the enumerations – this reduces the time complexity to O(*NL* log*L*).

**Figure 3:**
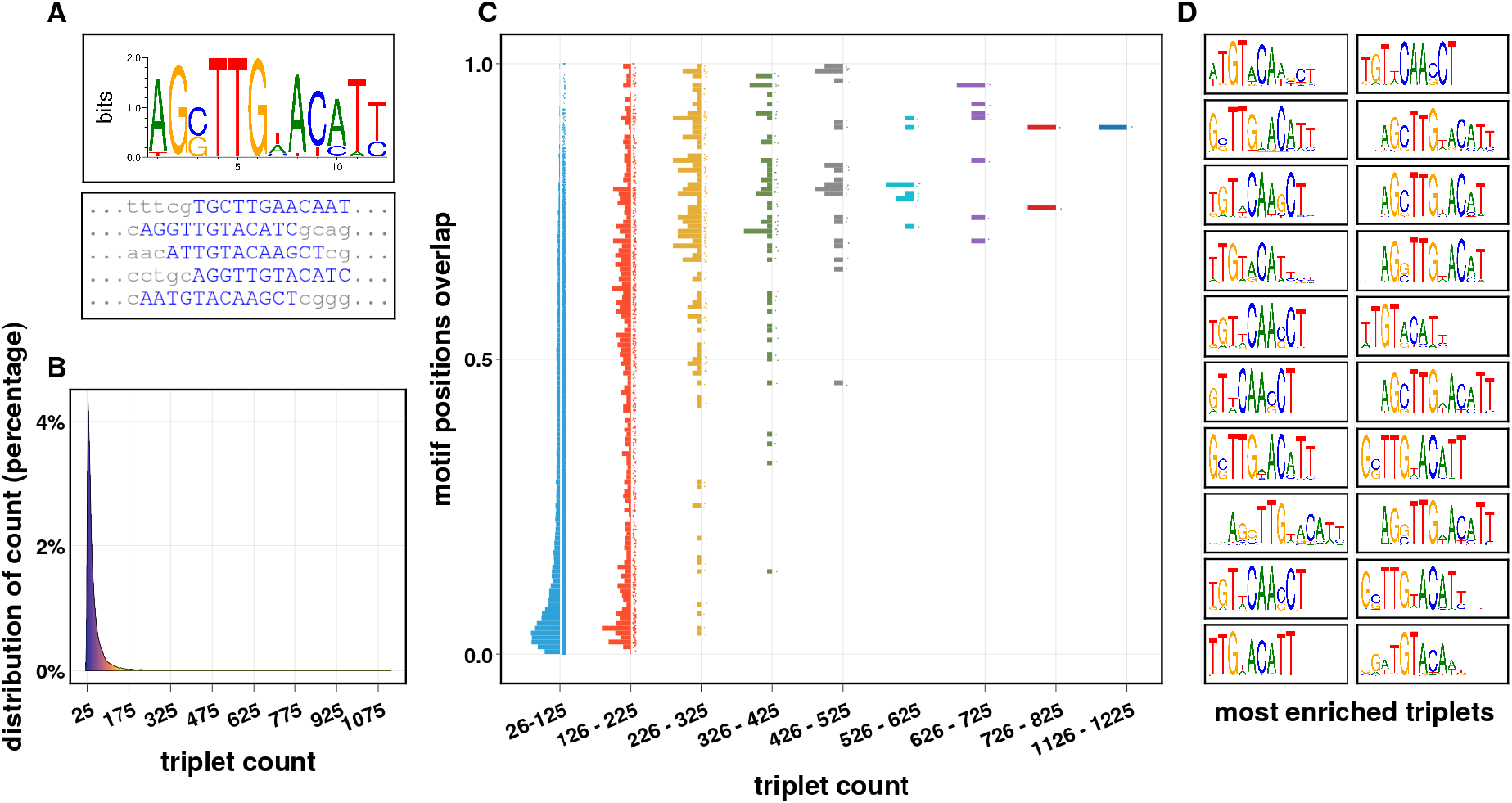
Illustration of the effectiveness of triplets on simple simulated data, using all sparse code components above the percentile *α* = 99.5% (equation 8). **A**. A ground truth motif and its realizations, in blue, are planted in 2,500 i.i.d. background strings (single motif, one occurrence per sequence, and equiprobable to occur in both forward and reverse complement direction). **B**. The empirical distribution of the triplet count, |***E***_*k*_ (·)|, for triplets with counts greater than 25. **C**. Distribution of motif position overlap for different ranges of the triplet counts. The motif position overlap is |*G* ∩ *T* |*/*|*T* |, where *G* is the positions (blue strings in A) of the planted motif and *T* is the positions covered by the triplet. This plot shows an upward trend of the overlap: the higher the triplet count, the more likely the set of positions covered by the triplet intersects the set of planted motif positions. **D**. The top 20 triplets with the highest counts, with the substrings they covered each turned into an MSA, and here visualized as PWMs.

**Figure 4:**
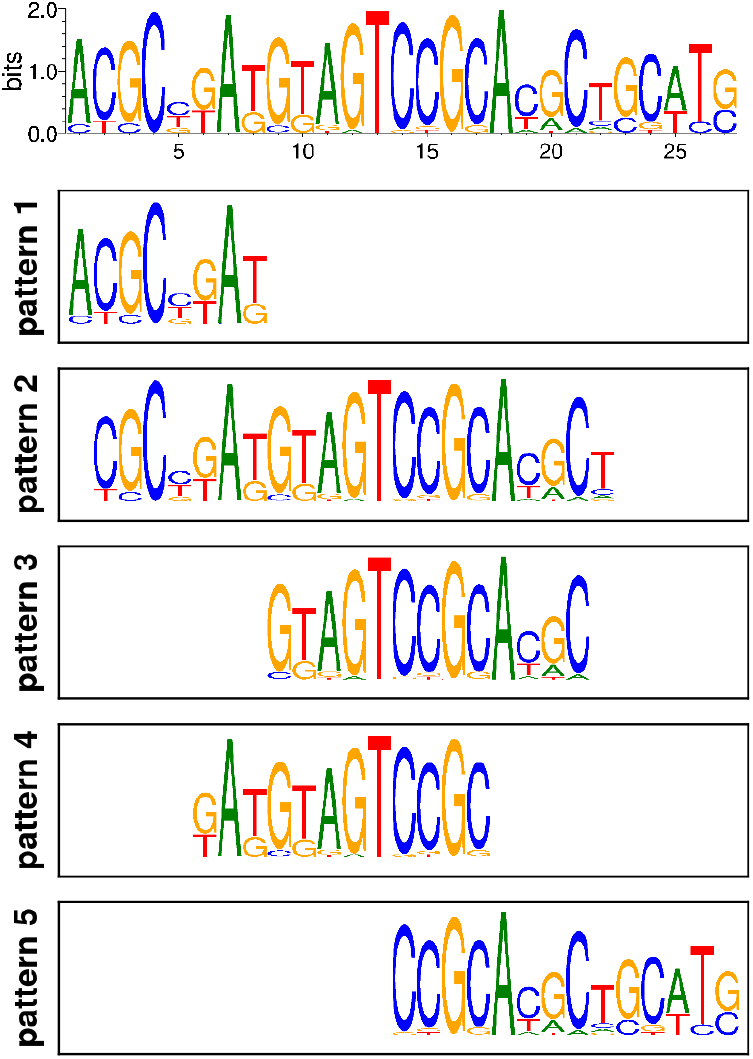
(example) Simulated ground truth motifs with five binding patterns, viewed as PWMs. The top PWM, not directly used in the simulated data, characterizes the intrinsic binding specificity of an imaginary protein, e.g., a Zinc finger protein. Each column is a point drawn from the Dirichlet distribution with a concentration of 0.1. We plant the instances from the five distinct binding patterns into the background strings.

**Figure 5:**
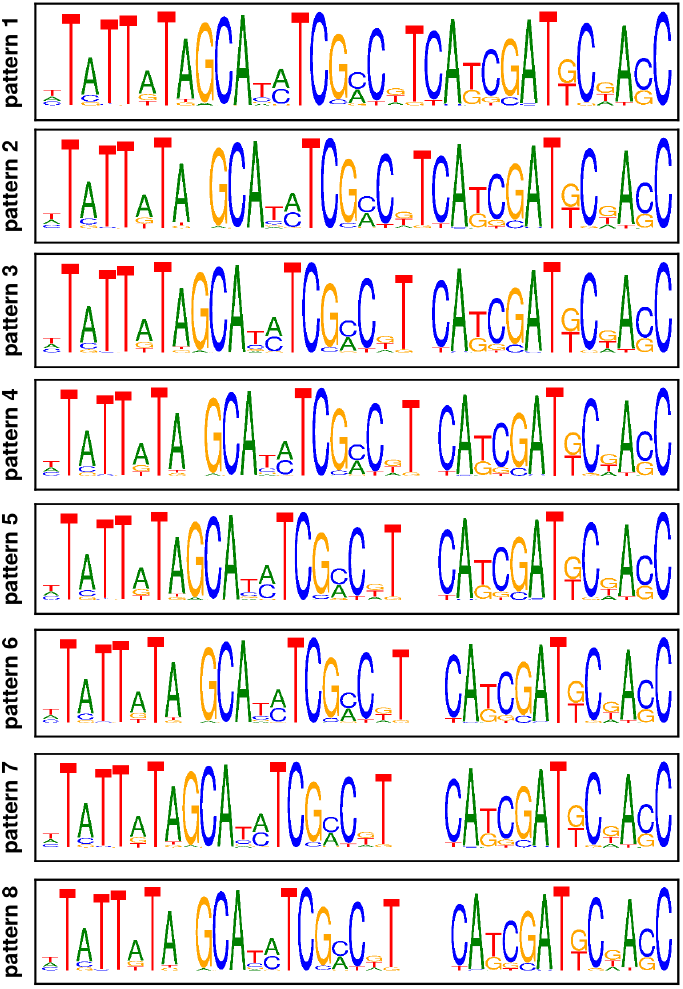
lated ground blocks with (example) A simutruth motif of three variable spacing. Two spacers are random variables *v*_12_∼*DiscreteUniform*(0, 1) and *v*_23_∼ *DiscreteUniform*(0, 1, 2, 3)that characterize the number of background nucleotide in between the adjacent blocks.

**Figure 6:**
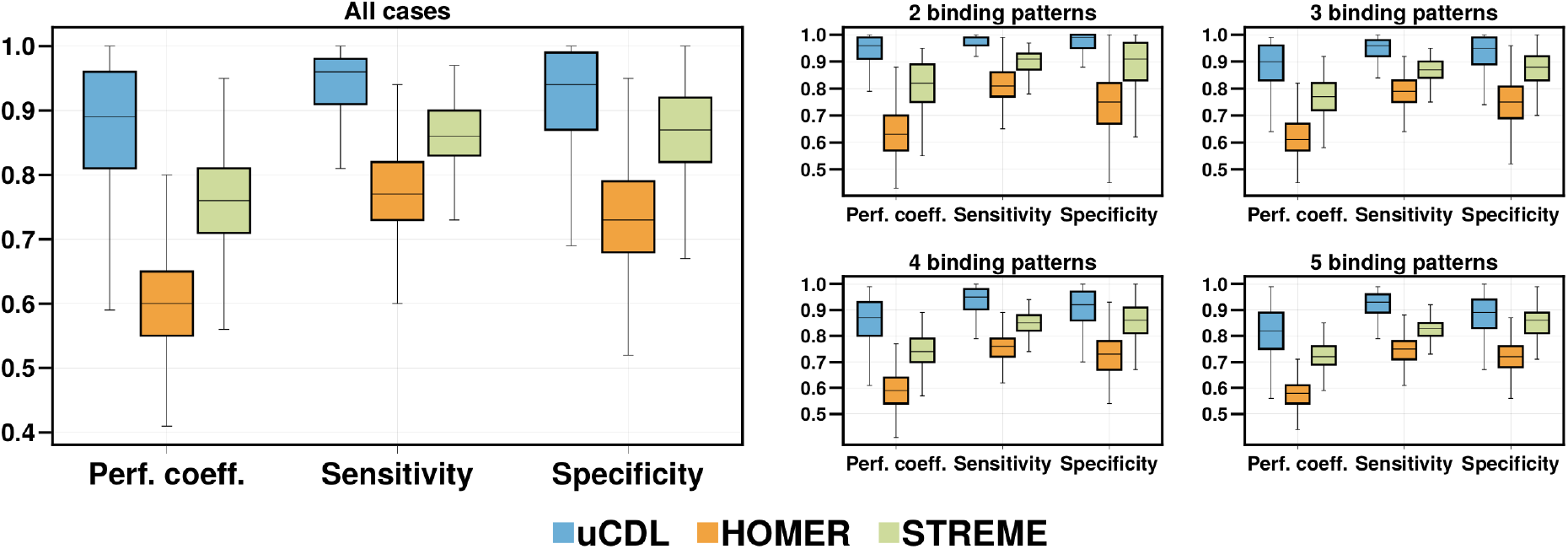
Performance on the simulated motifs with multiple binding patterns. The overall performance on all 2,000 simulated datasets is on the left, and separate cases in the 2,000 datasets with a fixed number of binding patterns are on the right. The midline in the interquartile range is the median. Our result shows that, for relatively conserved motifs in i.i.d. genomic background, uCDL on average is more sensitive and specific in all cases where multiple binding patterns exist, each is equiprobable to occur, and may share similar patterns.

**Figure 7:**
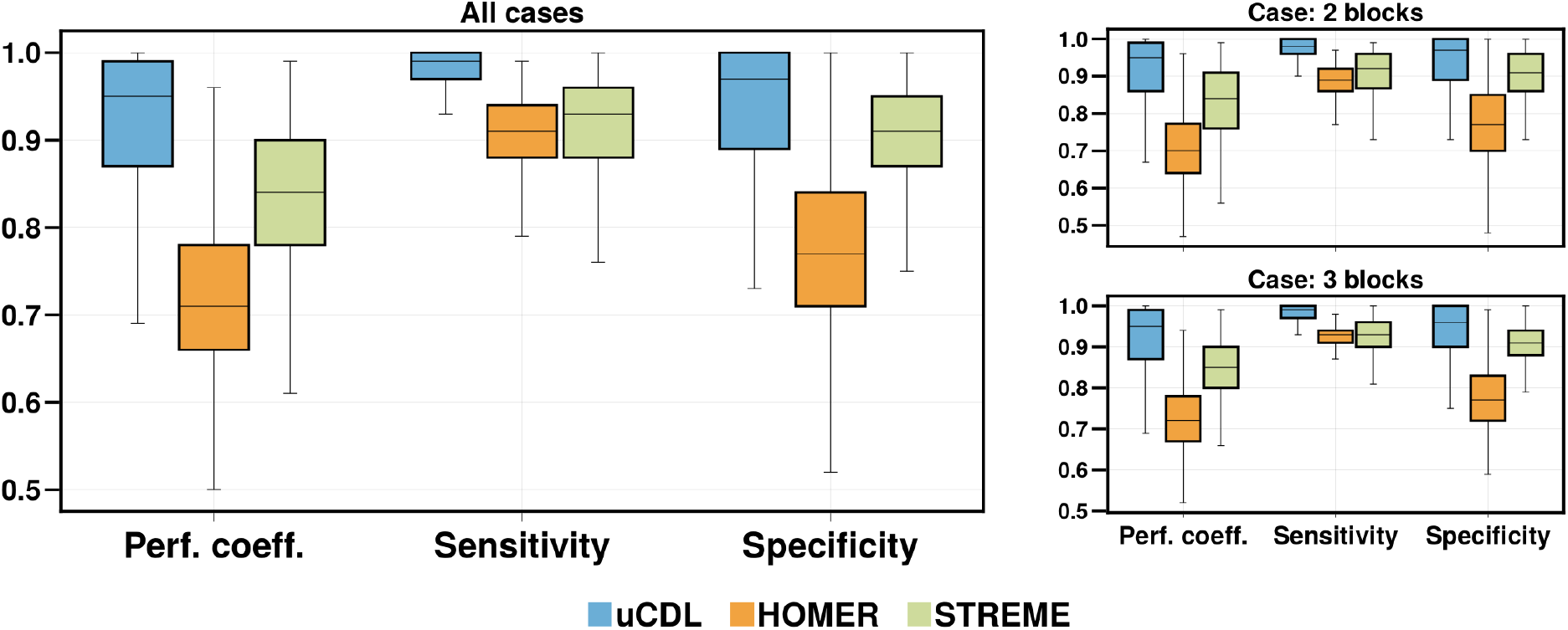
Performance on the simulated motifs with variable spacings. The overall performance on all 2,000 simulated datasets is on the left, and separate cases in the 2,000 cases with two blocks or three blocks of motifs are on the right. The midline in the interquartile range is the median. Our result shows that, for relatively conserved motifs in i.i.d. genomic background, uCDL is on average more sensitive and specific for two to three blocks of motifs with variable spacing.

We perform a generalized ESD test [Ros83] to identify the outliers in the triplet count distribution, which are the enriched triplets. With a suitable significance level and an upper bound to cap the number of outliers, we identify the enriched triplets and transform them into PWMs. Following the approach in [WS03], we merge similar PWMs using the average log-likelihood ratio. We expand/trim the width of PWMs by calculating the information content of adjacent positions/columns [SS90]. We then perform greedy alignment [SH89; HHS90; LMW99] with the merged PWMs ***P***_1_, …, ***P***_*J*_ to capture the binding sites the enriched triplets may not cover.

In the greedy alignment, each PWM ***P***_*j*_ is re-estimated with the aligned positions that each score above a threshold *τ*_*j*_. This step allows zero or more occurrences per sequence for each PWM. We obtain the scorethreshold *τ*_*j*_ from an approximation algorithm, pvalue2Score, with a well-chosen p-value [TV07]. If some positions are selected by more than one PWM, we let the PWM that achieves the highest score re-estimate with such positions. The greedy alignment ensures all the estimated PWMs are from sets of mutually disjoint positions. We use Fisher’s exact test on the held-out sequences to evaluate the significance of discovered motifs.

## 3 Results

We name our method as *unfolded convolutional dictionary learning* (uCDL). The following sections show results in both simulated and real biological data. For all experiments, we compare with STREME version 5.5.0 and HOMER version 4.11. The command line arguments used are streme --maxw 30 -p <input> -o <output> and findMotifs.pl <input> fasta <output>.

### 3.1 Simulated data: motifs with multiple binding patterns

To capture motifs of multimeric binding and multiple binding domains [SG14], we construct each of the simulated datasets in the following manner. The ground truth motifs are PFMs. Specifically, we draw a point from the Dirichlet distribution with a concentration parameter of 0.1 as each column in the ground truth PFM. We then

1. Determine the number of *binding patterns*, i.e., the number of motifs in the dataset, by drawing a number from *DiscreteUniform*(2,3,4,5).
2. Determine the width of each motif by drawing instances from *DiscreteUniform*(8, 9, …, 22).
3. Once the first motif ***G***_1_ is constructed with width *w*_1_, the second motif is constructed by drawing an overlap ratio *o* from ***o*** ∼ *Uniform*([0.1, 0.9]). We force the first columns of the second motif ***G***_2_ to be identical to columns ***G***_1_[:, *w*_1_ − ⌈ *o* × *w*_1_⌉ +1:*w*_1_]. The upper-bound 0.9 and the lower-bound 0.1 for the range of the random variable ***o*** guarantees that there is at least some overlap, but the two motifs are never identical.
4. If there are more than two binding patterns, as indicated by 1), the 3rd motif uniformly and randomly picks either the first or the second motif to overlap. Then we construct it following the procedure in 3). We generate the 4th and the 5th motif similarly if needed.
5. We generate 3,000 100bp sequences {***s***_*n*_} with i.i.d. background from *DiscreteUniform* {A,C,G,T}. For each sequence ***s***_*n*_, we uniformly and randomly chose a location to plant a motif instance, where we draw the instance from one of the ground truth motifs uniformly at random. The planted instance is equally likely to occur in its forward or reverse complement direction.

### 3.2 Simulated data: motifs with variable spacing

We construct each simulated dataset that contains motifs with variable spacing as follows:

1. Determine the binding sites as “blocks” of motifs by uniformly randomly selecting the number of blocks from *Uniform(2,3)*. Simulate the motif for each “block” in the same manner as step 2 in 3.1.
2. In between every two *i*-th and (*i*+1)-th blocks of motifs, we set a random variable *v*_*i,i*+1_ ∼ *DiscreteUniform*(1, *ν*) where *ν* is a number drawn from *DiscreteUniform*(1,10). The random variable *v*_*i,i*+1_ characterizes the variable spacing in between each adjacent block of motifs, and *ν* is the maximal number of nucleotides that separate the *i*-th and (*i*+1)-th motifs.
3. We generate 3,000 100bp sequences {***s***_*n*_} with i.i.d. background from *DiscreteUniform* {A,C,G,T}. For each sequence ***s***_*n*_, we uniformly and randomly chose a location to plant an instance of the ground truth motifs; we draw a motif instance from each block, and each *i*-th and (*i*+1)-th adjacent block is separated by the number of background nucleotides determined by drawing a number from each random variable *v*_*i,i*+1_’s. The planted instance is equally likely to occur in its forward or reverse complement direction

We generate 2,000 simulated datasets following the above procedure for each simulated data scenario: motifs with multiple binding patterns and motifs with variable spacing. For each simulated dataset, denote *G*_*i*_ and *D*_*j*_, the set of positions of the *i*-th ground truth motif and the *j*-th discovered motif, respectively. Let *G* = ∪_*i*_*G*_*i*_ be the union of the planted positions of all the ground truth motifs, and *D* = ∪_*j*_*D*_*j*_ be the union of positions of all the occurrences of the discovered motifs. The positions of occurrences of the discovered motifs are the positions that motifs (PWMs) scored above a threshold; we set each discovered PWM’s threshold using pvalue2score with p-value set to 2.5e-4 ([TV07]). Following [PS+00], we define the performance coefficient as |*G*∩*D*| */* |*G*∪*D*|, sensitivity |*G*∩*D*| */* |*D*|, and specificity |*G*∩*D*| */* |*G*| as the performance measure. We calculated the performance measures on the test sets: for each simulated dataset, the test set is another newly generated 3,000 100bp strings using its respective ground truth.

### 3.3 Biological data

An example that highlights the difference between motifs discovered by uCDL compared to STREME and HOMER is in figure 8. In figure 8, all methods find the reference motif from JASPAR. Additionally, uCDL finds an alternative binding pattern (uCDL-3) absent from STREME and HOMER’s results — a pattern CTTCC appended to the left of the reference motif, and a long motif (uCDL-4) that corresponds to the L1PA5 transposable element that is known to bind the YY1 transcription factor [ABM04; Sch+13]. The positions 10 to 19 of uCDL-4 overlap the reference motif. The long motif uCDL-4 is only partially discovered by STREME (STREME-2 and STREME-3, which corresponds to positions 23 and 41 of uCDL-4 in figure 8 part **(E)**. We suspect heuristic such as substring-masking leads to patterns such as uCDL-3 not discovered by STREME and HOMER.

**Figure 8:**
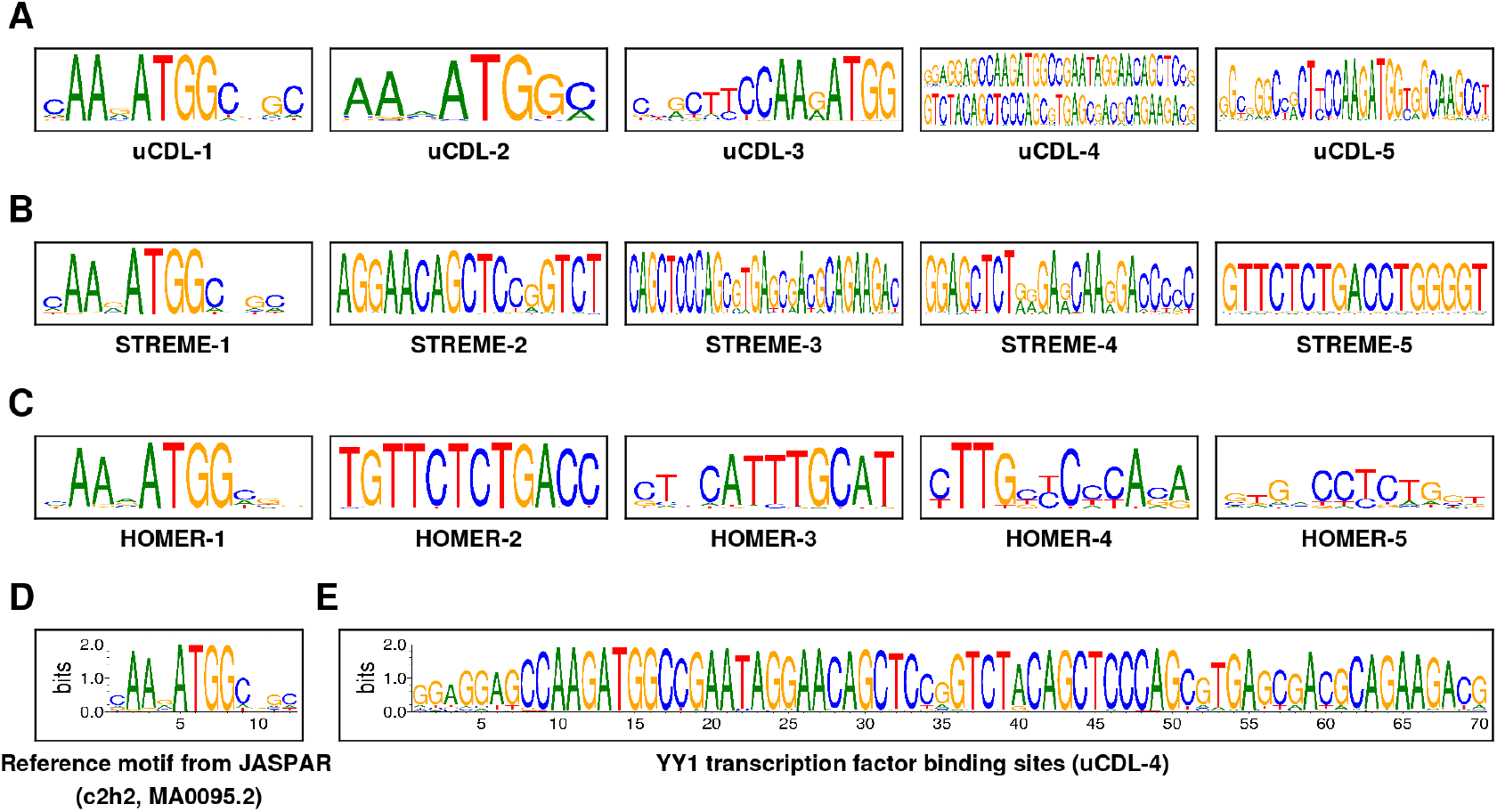
Motifs discovery results on MA0095.2 (C2H2) from JASPAR [Cas+22]. We list the significant reported motifs from each software in the order of statistical significance. We have the top 5 motifs **A**. discovered by uCDL. **B**. discovered by STREME. **C**. discovered by HOMER. **D**. Reference motif from JASPAR. **E**. The L1PA5 transposable element that is known to bind the YY1 transcription factor [ABM04; Sch+13], rediscovered by uCDL.

Many datasets contain motifs that co-occur in the same sequence, especially when the two motifs are identical. We list some of such instances from the JASPAR datasets in figure 9. Our observations correspond to several modes of DNA binding pointed out in previous studies [SG14]. For example, the datasets from the BZIP family (MA0462.1, MA0478.1) contain adjacent TGAC half-sites that are concentrated at around 12 or 40 nucleotides apart from each other. The motifs in the nuclear receptors datasets (MA0114.2, MA0505.1, MA0512.1, not shown in figure) also show that different length spacers separate the AGGTCA half-sites.

**Figure 9:**
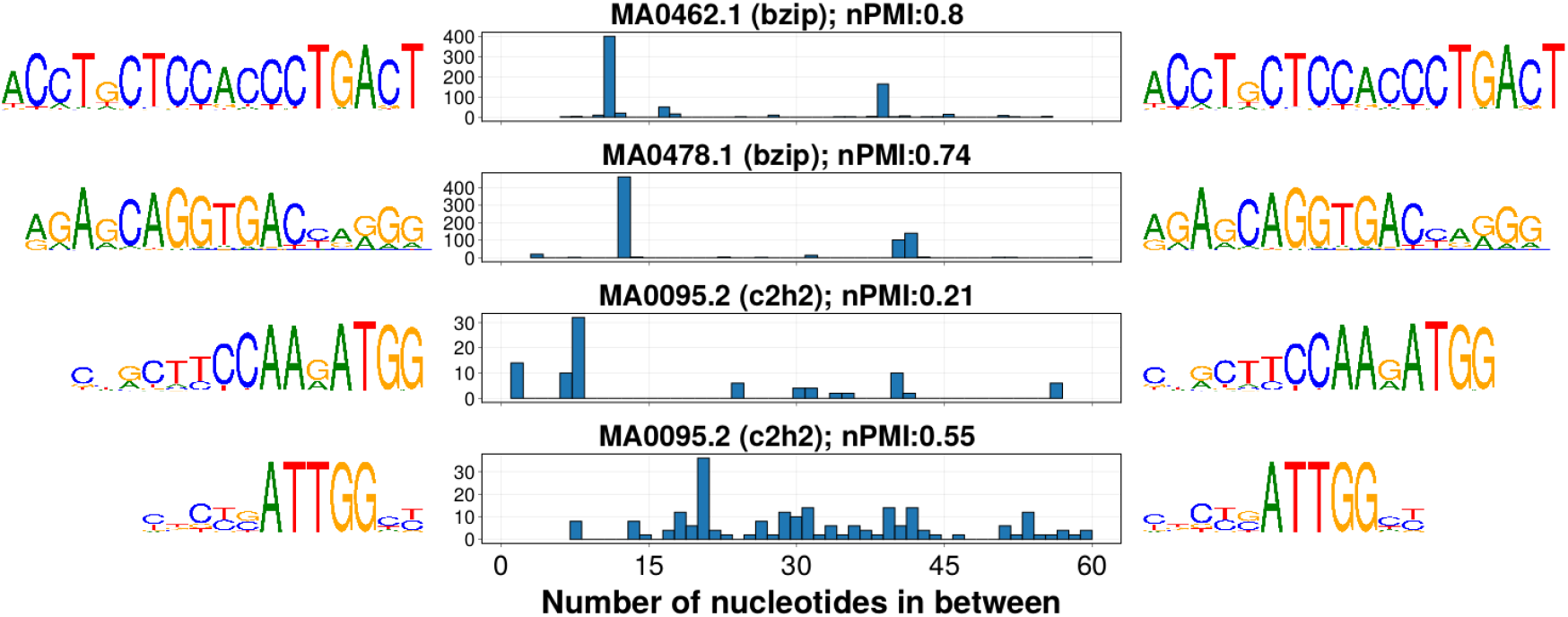
Motif co-occurrence on the same DNA string from some JASPAR datasets [Cas+22], not reported on JASPAR. All the motifs are statistically significant (Fisher’s exact test, *p <* 1e − 6). The normalized pairwise mutual information (nPMI) measures the degree of co-occurrence, where a value of 0 indicates no co-occurrence and a value of 1 indicates complete co-occurrence (Supplementary file 1, section C.2). The histogram considers all the co-occurring pairs on the same string, including the motifs that may co-occur more than once on the same string.

Some datasets contain motifs that exhibit multimeric binding [SG14], i.e., a core motif and other motifs that are the expanded version of the core motif may exist. We observe such binding mode in the BZIP, C2H2, p53 proteins, and others (See supplementary file 2). Some datasets contain motifs with multiple DNA binding domains, e.g., we find the dataset MA0162.2 from JASPAR has another significant motif consisting of consecutive 2mers of C. Some datasets from JASPAR contain significant motifs that show alternative structural conformation, e.g., MA0102.3. We show these results in figure 10. Note that the binding mode of a protein is not mutually exclusive. For example, the significant motifs in the dataset MA0095.2 from JASPAR suggest that a protein can have binding modes exhibiting variable spacing and multimeric binding.

**Figure 10:**
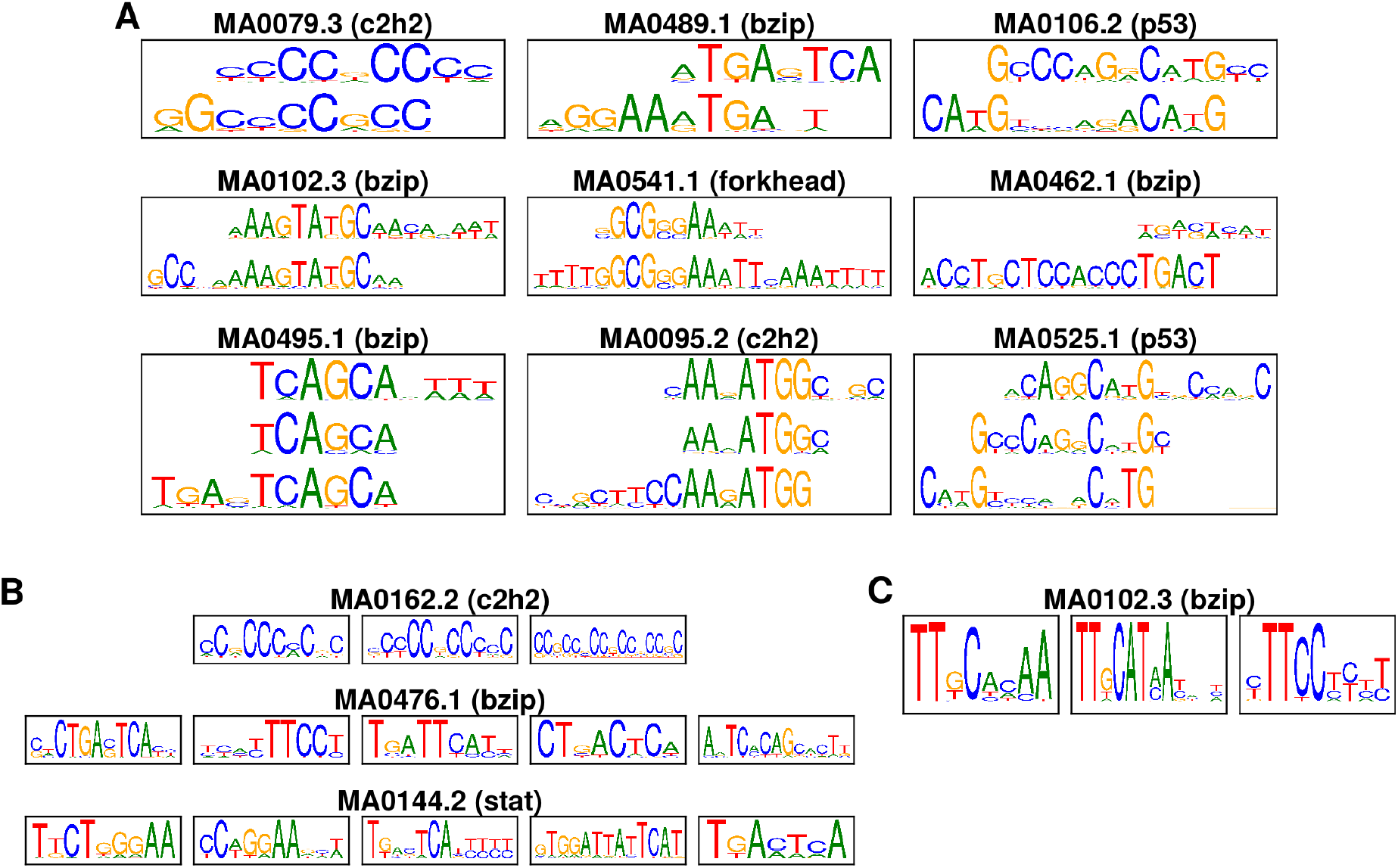
Motifs with **A**. multimeric binding **B**. multiple DNA binding domains **C**. alternate structural conformation [SG14] from some JASPAR datasets [Cas+22], not reported on JASPAR. All the motifs are statistically significant (Fisher’s exact test, *p <* 1e−6).

We also found that long motifs exist in some datasets from JASPAR. These long motifs may be transposable elements and often contain the core motif as its sites. We show some of the long motifs we discovered in figure 11.

**Figure 11:**
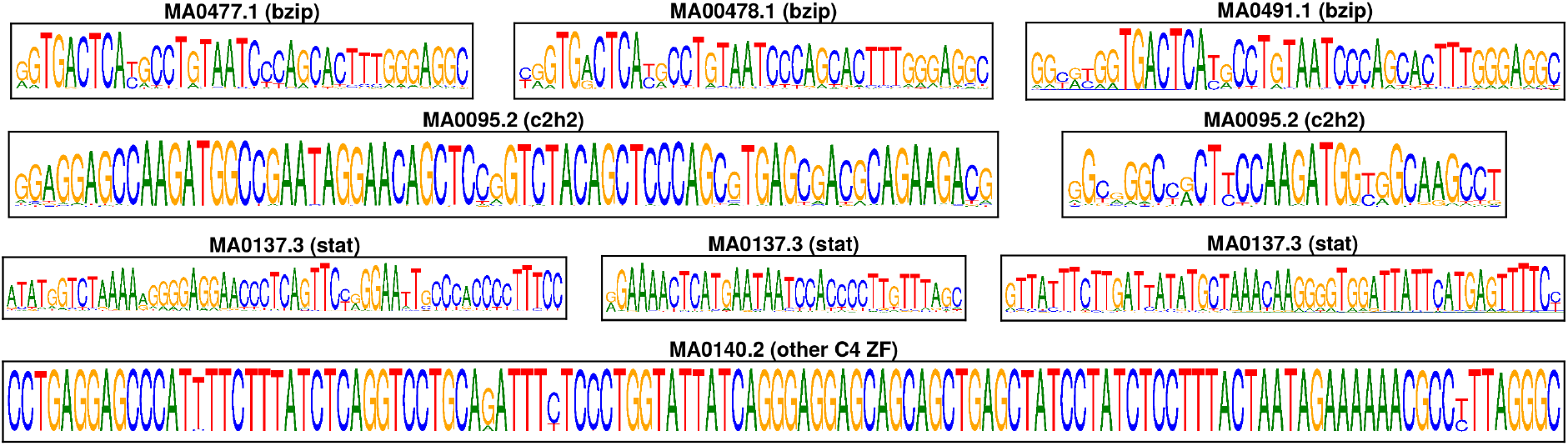
Some long motifs discovered from the JASPAR datasets [Kha+18], not reported on JASPAR. All the motifs are statistically significant (Fisher’s exact test, *p <* 1e−6)

### 3.4 Speed

uCDL finds motifs for large datasets in a reasonable time. As shown in figure 12, using simulated data in section 3.1, it takes on average about 13 minutes to find motifs for a dataset that contains 60,000 100bp input DNA strings. Figure 12 shows that the runtime of uCDL is linear to the input size. We performed the tests on a computer with an NVidia Geforce RTX 3090 GPU, AMD Ryzen 5 5600X 6-Core Processor, and 64 GB of RAM. The CDL model consists of 160 length-8 filters, and we trained the network for 25 epochs. Further improvement is possible, for example, if we could design a learning-based technique that learns a map to replace the approximation algorithm pvalue2score [TV07] for the PWM score threshold.

**Figure 12:**
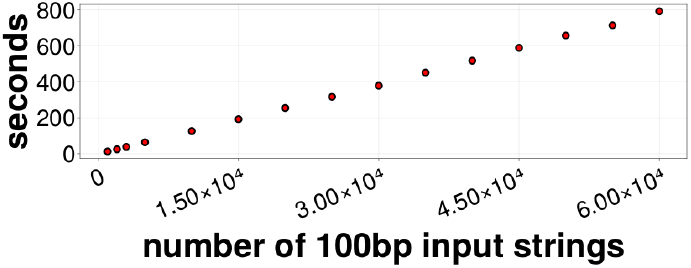
Average run of 10 simulated datasets (section 3.1) for motif discovery with uCDL.

## 4 Discussion

Some may ask, as we set the filters as position frequency matrix in section 2.2: why do we not interpret the filters as the motifs? The answer is that since our model is in a distributed representation, it rarely represents a substring using a single filter. The filters are almost always jointly, i.e., linearly combined to represent the substrings. Further, the batch size used to train the unfolded CDL (uCDL) network can also affect how each filter is learned; we show this in supplementary file 1, section B.3. Our results show that the triplet (section 2.5) is a more reliable way to look at frequently occurring patterns. We constrain the filters as position frequency matrices simply because this is more interpretable than just an ordinary feature vector.

The uCDL (section 2.4) is a different model than the CDL model (section 2.2) but enjoys some remarkable advantages over the CDL without sacrificing model interpretability. An advantage of uCDL is that the training is much faster than CDL – we no longer compute the gradient using all data points as in typical optimization subroutines – we now train the model using backpropagation with stochastic gradient descent. As a model, uCDL is a neural network with a structural bias to learn a sparse representation. Every layer in the uCDL has a clear interpretation, as it mimics the alternating minimization of CDL, outlined in section 2.3. Another advantage of uCDL is that it is more expressive than the CDL model, e.g., the sparsity-inducing parameter *λ* and the step-sizes *η*_*t*_ and *μ* in CDL are now parameters of the network, no longer require hyperparameter tuning.

A limitation of uCDL is that we do not obtain a unique sparse representation for each run of the motif discovery. Motif discovery methods do not generally provide identifiability guarantees, especially for overparameterized models such as neural networks. However, our implementation of uCDL will only report statistically significant motifs, which should be mostly identical in each run. We encourage users who use our program to train uCDL multiple times when in doubt.

We can use the computational graph in figure 2 for various downstream regulatory genomics problems such as functional DNA classifications ([Gha+14]) and promoter strength estimations ([Jor+22]). We can view the computational graph as an encoder-decoder architecture: the encoder takes the DNA strings as input and outputs a sparse code. We can then design a decoder that takes the sparse code for other applications. Attacking regulatory genomics problems from such an approach provides advantages for interpretability as the sparse code carries syntactic and semantic structures in the DNA strings. The filters can be viewd as “soft” kmers and that sparse coding is a trick for finding kmers enrichment without explicit enumerations.

## 5 Usage

Our method written in Julia [Bez+17], and is available under the MIT license as open source software at https://github.com/kchu25/UnfoldCDL.jl.

## 6 Conclusion

We present our unfolded convolutional dictionary learning (uCDL) method for motif discovery. Both simulated and experimental data suggest that uCDL is more effective at capturing motifs that exhibit complex binding modes than methods such as STREME and HOMER. The complex binding modes include variable spacing, multimeric binding, multiple DNA binding domains, and alternative structural conformation [SG14]. Our method is the first that infers the motifs via a sparse representation, a distributed representation model with intuitive built-in interpretations that allow for efficient inference. Our approach is scalable to large datasets. We can extend the resulting computational graph from deep unfolding for downstream regulatory genomics problems to extract biological strings’ syntactic and semantic structures.

## 7 Acknowledgement

We thank Ting Wang for help identifying the L1 repetitive element and Ulugbek Kamilov for the suggestion of using deep unfolding to scale up convolutional dictionary learning. This research is funded by NIH grant 1R01GM125736 and internal funds from Washington University School of Medicine.

## Footnotes

1 In practice, since we are doing 1D convolutions, we have that _***X***_ ≜ ***x***_*n*_ : ***x***_*mn*_ **0, *P x***_*mn*_ = **0**, where ***P*** is a projector that maps each equally spaced three-consecutive components of the sparse code to zero.

## Notes

https://jaspar.genereg.net/

https://github.com/kchu25/UnfoldCDL.jl

### Competing Interest Statement

The authors have declared no competing interest.

### Summary of Updates

Update to the correct github link in the manuscript.

## References

[NY83] Arkadij Semenovič Nemirovskij and David Borisovich Yudin. “Problem complexity and method efficiency in optimization”. In: (1983).

[Ros83] Bernard Rosner. “Percentage points for a generalized ESD many-outlier procedure”. In: Technometrics 25.2 (1983), pp. 165–172.

[Hin84] Geoffrey E Hinton. “Distributed representations”. In: (1984).

[SH89] Gary D Stormo and George W Hartzell. “Identifying protein-binding sites from unaligned DNA fragments”. In: Proceedings of the National Academy of Sciences 86.4 (1989), pp. 1183–1187.

[HHS90] Gerald Z Hertz, George W Hartzell Iii, and Gary D Stormo. “Identification of consensus patterns in unaligned DNA sequences known to be functionally related”. In: Bioinformatics 6.2 (1990), pp. 81–92.

[SS90] Thomas D Schneider and R Michael Stephens. “Sequence logos: a new way to display consensus sequences”. In: Nucleic acids research 18.20 (1990), pp. 6097–6100.

[BE95] Timothy L Bailey and Charles Elkan. “Unsupervised learning of multiple motifs in biopolymers using expectation maximization”. In: Machine learning 21.1 (1995), pp. 51–80.

[LMW99] Ming Li, Bin Ma, and Lusheng Wang. “Finding similar regions in many strings”. In: Proceedings of the thirty-first annual ACM symposium on Theory of computing. 1999, pp. 473–482.

[PS+00] Pavel A Pevzner, Sing-Hoi Sze, et al. “Combinatorial approaches to finding subtle signals in DNA sequences.” In: ISMB. Vol. 8. 2000, pp. 269–278.

[BT03] Amir Beck and Marc Teboulle. “Mirror descent and nonlinear projected subgradient methods for convex optimization”. In: Operations Research Letters 31.3 (2003), pp. 167–175.

[WS03] Ting Wang and Gary D Stormo. “Combining phylogenetic data with co-regulated genes to identify regulatory motifs”. In: Bioinformatics 19.18 (2003), pp. 2369–2380.

[ABM04] Jyoti N Athanikar, Richard M Badge, and John V Moran. “A YY1-binding site is required for accurate human LINE-1 transcription initiation”. In: Nucleic acids research 32.13 (2004), pp. 3846– 3855.

[TV07] Hélène Touzet and Jean-Stéphane Varré. “Efficient and accurate P-value computation for Position Weight Matrices”. In: Algorithms for Molecular Biology 2.1 (2007), pp. 1–12.

[GL10] Karol Gregor and Yann LeCun. “Learning fast approximations of sparse coding”. In: Proceedings of the 27th international conference on international conference on machine learning. 2010, pp. 399–406.

[BEL13] Hilton Bristow, Anders Eriksson, and Simon Lucey. “Fast convolutional sparse coding”. In: Proceedings of the IEEE Conference on Computer Vision and Pattern Recognition. 2013, pp. 391–398.

[Sch+13] Petra C Schwalie et al. “Co-binding by YY1 identifies the transcriptionally active, highly conserved set of CTCF-bound regions in primate genomes”. In: Genome biology 14.12 (2013), pp. 1– 15.

[Gha+14] Mahmoud Ghandi et al. “Enhanced regulatory sequence prediction using gapped k-mer features”. In: PLoS computational biology 10.7 (2014), e1003711.

[SG14] Trevor Siggers and Raluca Gordân. “Protein–DNA binding: complexities and multi-protein codes”. In: Nucleic acids research 42.4 (2014), pp. 2099–2111.

[HHW15] Felix Heide, Wolfgang Heidrich, and Gordon Wetzstein. “Fast and Flexible Convolutional Sparse Coding”. In: Proceedings of the IEEE Conference on Computer Vision and Pattern Recognition (CVPR). June 2015.

[Bez+17] Jeff Bezanson et al. “Julia: A fresh approach to numerical computing”. In: SIAM review 59.1 (2017), pp. 65–98.

[LL17] Scott M Lundberg and Su-In Lee. “A unified approach to interpreting model predictions”. In: Advances in neural information processing systems 30 (2017).

[Woh17] Brendt Wohlberg. “Convolutional sparse coding with overlapping group norms”. In: arXiv preprint arXiv:1708.09038 (2017).

[DI18] Bogdan Dumitrescu and Paul Irofti. Dictionary learning algorithms and applications. Springer, 2018.

[GW18] Cristina Garcia-Cardona and Brendt Wohlberg. “Convolutional dictionary learning: A comparative review and new algorithms”. In: IEEE Transactions on Computational Imaging 4.3 (2018), pp. 366–381.

[Kha+18] Aziz Khan et al. “JASPAR 2018: update of the open-access database of transcription factor binding profiles and its web framework”. In: Nucleic acids research 46.D1 (2018), pp. D260–D266.

[KP21] Peter K Koo and Matt Ploenzke. “Improving representations of genomic sequence motifs in convolutional networks with exponential activations”. In: Nature machine intelligence 3.3 (2021), pp. 258–266.

[MLE21] Vishal Monga, Yuelong Li, and Yonina C Eldar. “Algorithm unrolling: Interpretable, efficient deep learning for signal and image processing”. In: IEEE Signal Processing Magazine 38.2 (2021), pp. 18–44.

[Cas+22] Jaime A Castro-Mondragon et al. “JASPAR 2022: the 9th release of the open-access database of transcription factor binding profiles”. In: Nucleic acids research 50.D1 (2022), pp. D165–D173.

[Jor+22] Tobias Jores et al. “Learning the Grammar of Regulatory DNA in Plants”. In: Plant and Animal Genome XXIX Conference (January 8-12, 2022). PAG.2022.

[Nov+22] Gherman Novakovsky et al. “Obtaining genetics insights from deep learning via explainable artificial intelligence”. In: Nature Reviews Genetics (2022), pp. 1–13.

