## Supplemental File 1 for "Deep unfolded convolutional dictionary learning for motif discovery"

### A Performance on other simulated data

We compare with STREME version $5.5.0$ and HOMER version $4.11$ for all the experiments conducted in this work. The command line arguments used are

```
streme --maxw 30 -p <input> -o <output>
```

for STREME, and

```
findMotifs.pl <input> fasta <output>
```

for HOMER.

For a fair performance comparison, we use only significant motifs (PWMs) found by STREME and HOMER (e-value less than $1$ for STREME and p-value less than $1$e-$12$, i.e., the ones that are labeled as "likely false positive" for HOMER). Each column on both the left-end and the right-end of the returned PWMs is removed if the column has information content less than 0.01. Note that the information content [SS90] of the $j$-th column of PWM $\boldsymbol{P}$ with the uniform background assumption is defined as

$$2 + \sum_{a} \boldsymbol{P}[a,j]\log_2 \boldsymbol{P}[a,j]. \tag{1}$$

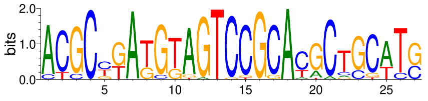

**Figure 1:** A simulated ground truth motif as a PWM; the genomic background is assumed to be uniform and each column is a point sampled from a Dirichlet distribution with a concentration of 0.1.

In all the experiments on simulated data, the simulated PFMs ground truths have columns that are points drawn from the Dirichlet distribution with the concentration parameter set to $0.1$. We show an example of such PFM instance as a PWM in figure 1.

For each dataset, denote $G_i$ and $D_j$, the set of positions of the $i$-th ground truth motif and the $j$-th discovered motif, respectively. Let $G = \cup_i G_i$ be the union of positions of all the ground truth motifs, and $D = \cup_j D_j$ be the union of positions of all the discovered motifs. Following Pevzner and Sze [PS+00], we use the following three quantities as the performance measure:

$$\text{performance coefficient: } \frac{|G \cap D|}{|G \cup D|} \qquad \text{sensitivity: } \frac{|G \cap D|}{|D|} \qquad \text{specificity: } \frac{|G \cap D|}{|G|} \tag{2}$$

#### A.1 Performance on motifs with single binding mode, OOPS and ZOOPS

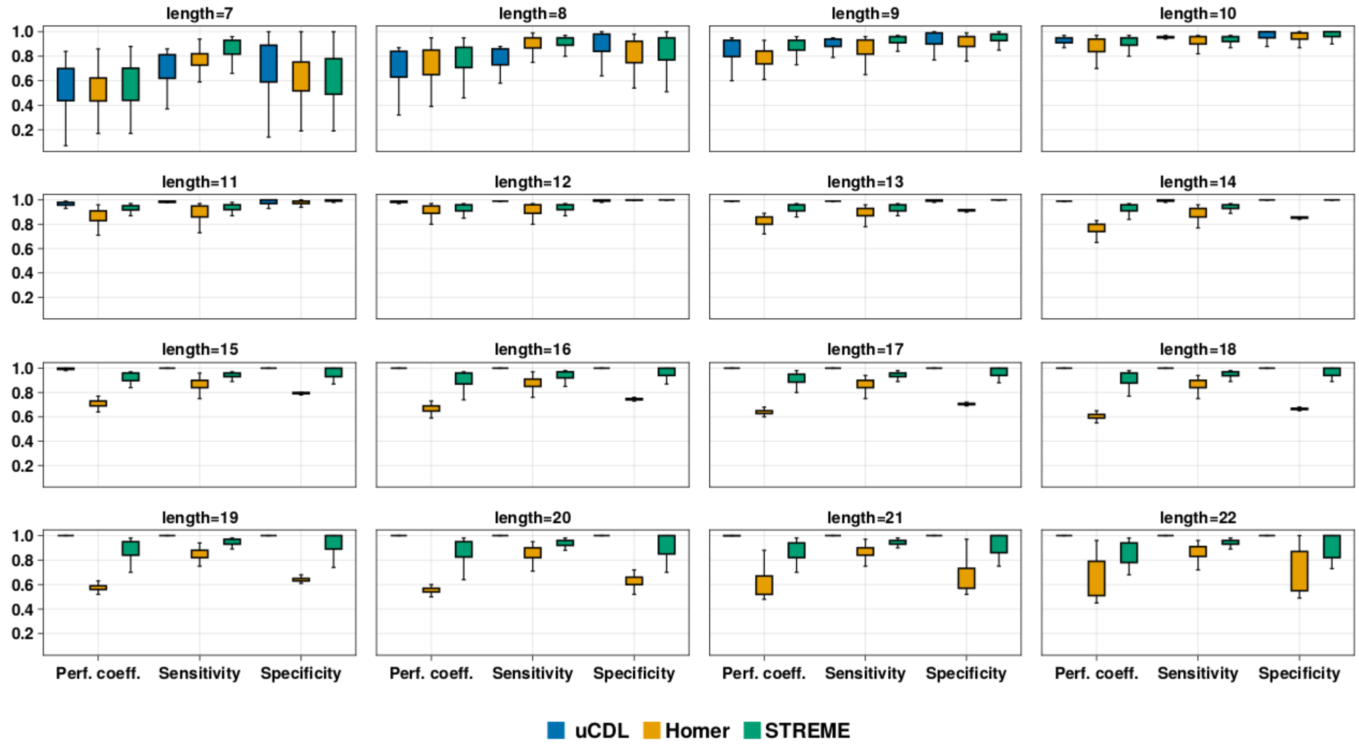

**Figure 2:** Performance measures on the simulated data of motifs with single binding mode with varying motif length. Each realization from the ground truth PFMs is planted at a random location in the background string uniformly at random. The background strings are generated in an i.i.d. fashion simply using a discrete uniform distribution on the alphabet { A,C,G,T }. Each dataset has one motif and one motif occurrence per string (OOPS). We test the results on another 100 newly generated datasets for each case. The length in each title indicates the motif length, i.e., how many nucleotides the motif spans. Our results show that all methods struggle when the length of the motif is small, as the short motifs are harder to be differentiated from the genomic background. HOMER by default performs less well for long motifs compared to CDLT and STREME.

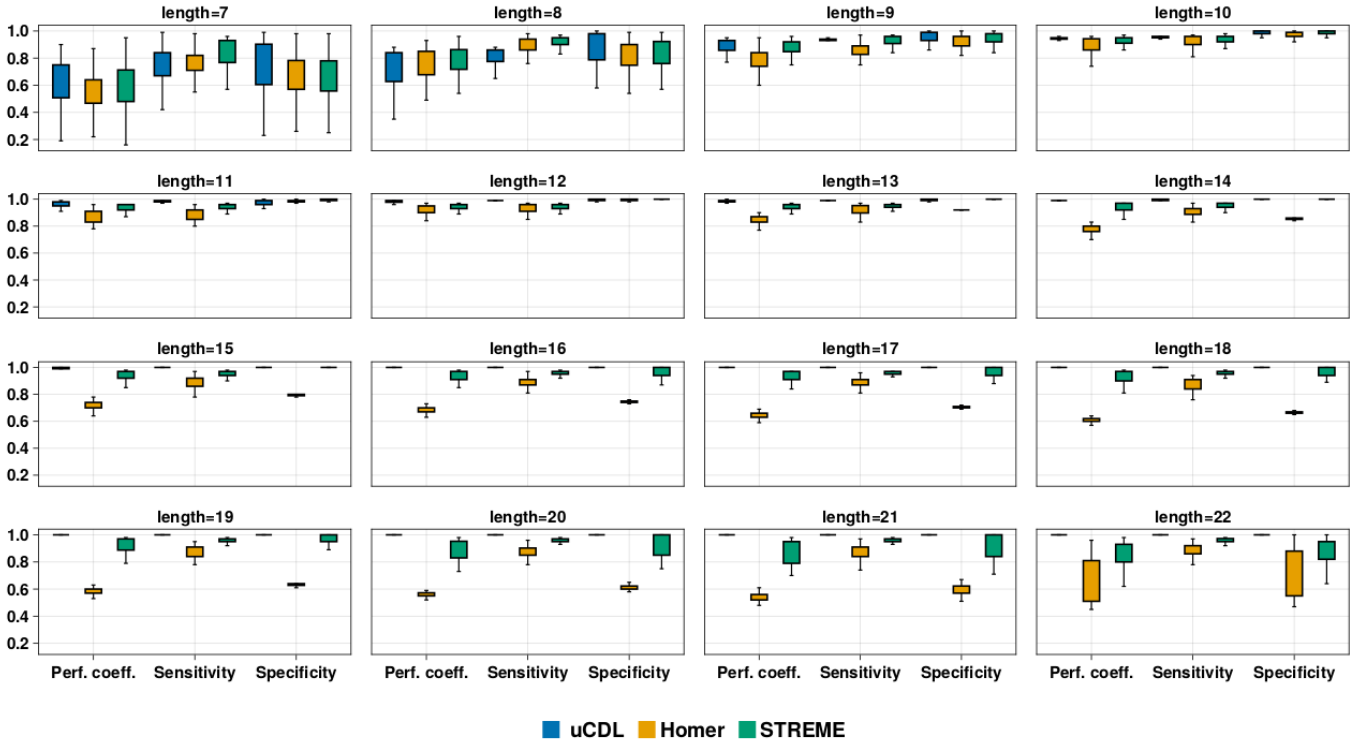

**Figure 3:** Same setting as figure 2, except that now each string in each dataset has zero or one motif occurrence (ZOOPS). We plant motifs in 60% of the strings in each dataset.

#### A.2 Performance on overlapping motifs, ZOOPS

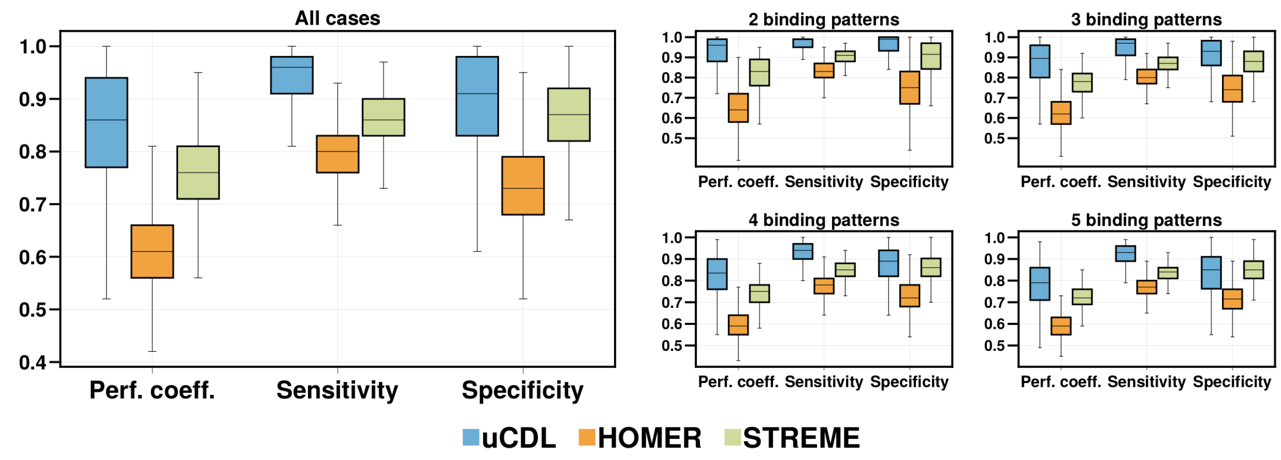

**Figure 4:** Performance comparison on simulated data on multiple binding patterns, using the same data generation shown in section 3.1, except that each input string now may contain zero motif. We plant motifs in 60% of the strings in each dataset.

#### A.3 Performance on motifs with variable spacings, ZOOPS

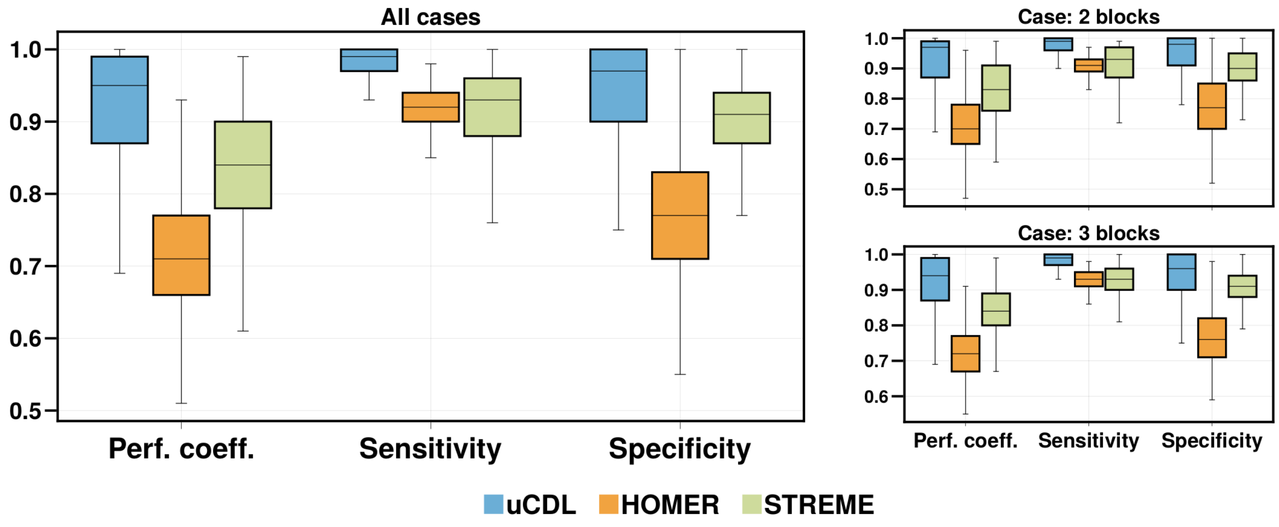

**Figure 5:** Performance comparison on simulated data on motifs with variable spacing, using the same data generation shown in section 3.2, except that each input string now may contain zero motif. We plant motifs in 60% of the strings in each dataset.

### B Implementation

#### B.1 Programming language and packages

We implement our method unfolded CDL (uCDL) in the Julia programming language [Bez+17], available from the Julia registry via the command `add uCDL`. We use Flux.jl and Zygote.jl [**innes2019differentiable**; Inn18] for the auto-differentiation on the computational graph in figure 2 in the main paper. We implement a customized kernel code for speeding up the greedy alignment using CUDA.jl [BFD18]. We render the PWM logo with Weblogo [Cro+04] and the rest of the visualizations with Makie.jl [DK21]. We generate all simulated data via Distributions.jl.

#### B.2 Hyperparameters for unfolded convolutional dictionary learning (uCDL) and other implementation details

##### B.2.1 Training the network

All the results shown in this paper for the CDL used the following hyperparameter setting: the number of filters is $M = 160$. We set each filter $\boldsymbol{d}_m$ to cover $8$-nucleotides. We train the unrolled CDL as a neural network with a batch size set to $48$ input strings; we carried out the backpropagation with Adabelief [Zhu+20]. The filter-diversity regularization parameter $\gamma$ in equation 7 is set to 5e-3. We perform four CSC iterations in the computational graph, and we add one iteration of the dictionary update (DU) after the CSC iterations. The reason for one iteration of DU is that the backpropagation of the network will also update the filters in the dictionary (since the dictionary is the network's weight). The primary reason for executing one iteration of DU in the forward pass is that it defines the data fitting loss and the interpretation of the learned filters.

##### B.2.2 Merge the PWMs derived from the triplets

Once we obtained the learned sparse representation, we find the enriched triplet by first constructing the set $\boldsymbol{V}_\alpha$ (equation 8) by setting $\alpha = 99.5\%$. Next, we use the generalized ESD test with a significance level set to $0.05$ and cap the number of outliers with an upper bound set to $1,000$ [Ros83]. The neighborhood parameter $\delta$ for the triplet is set to $8$ so that each triplet covers $8$ to $16$ nucleotides. We compare and merge PWMs derived from the triplets using the average log-likelihood ratio (ALLR) [WS03]. Note that given two PWMs $\boldsymbol{P}_1$ and $\boldsymbol{P}_2$, their corresponding count matrices $\boldsymbol{C}_1, \boldsymbol{C}_2$ and frequency matrices $\boldsymbol{F}_1, \boldsymbol{F}_2$, and the assumed genomic background $\boldsymbol{B}_\mathsf{A}, \boldsymbol{B}_\mathsf{C}, \boldsymbol{B}_\mathsf{G}, \boldsymbol{B}_\mathsf{T}$, the ALLR of column $i$ of $\boldsymbol{P}_1$ and column $j$ for $\boldsymbol{P}_2$ is defined as

$$\frac{\sum_\alpha \boldsymbol{C}[\alpha, i] \log_2 \left( \boldsymbol{F}[\alpha, j] / \boldsymbol{B}_\alpha \right) \; + \; \sum_\alpha \boldsymbol{C}[\alpha, j] \log_2 \left( \boldsymbol{F}[\alpha, i] / \boldsymbol{B}_\alpha \right)}{\sum_\alpha \boldsymbol{C}[\alpha, i] + \boldsymbol{C}[\alpha, j]}. \tag{3}$$

The physical interpretation of ALLR is that it measures how two proteins binding to each other's binding sites [WS03]. A "good" ALLR value implies that two proteins accept each other's sites or that their sites are equivalent. We merge pairs of PWMs derived from the triplets as long as there exists an alignment such that the average ALLR of that alignment of two PWMs is more than $0.25$ and that the two PWMs have small length differences (less than $4$). We merge such a pair by adding the corresponding two count matrices according to that alignment; if one PWM is shorter than the other, then we go back to the dataset and reconstruct the missing columns, using all the substrings that are still "inside" the dataset.

##### B.2.3 Expand/Trim the merged PWMs

Once we merge the PWMs derived from the triplets, we obtain a set of merged PWMs. We next expand the width of the merged PWM if each merged PWM's respective MSA has neighboring adjacent columns that are not included in the MSA and have information content higher than $0.7$ bits. If such neighboring columns do not exist, we "trim" the width of a PWM of both ends if either end of the PWM has a column that has information content lower than $0.4$ bits. We find such information content thresholds give good empirical performances.

###### B.2.4 Greedy alignment

We perform greedy alignment using the merged PWMs that have been expanded or trimmed. We say a substring is a "binding site" of each merged PWM $\boldsymbol{P}_j$ if $\boldsymbol{P}_j$ has scored above a score-threshold $\tau_j$ to that substring. We determine the score threshold $\tau_j$ of each $\boldsymbol{P}_j$ by the approximation algorithm pvalue2score with input p-value 2.5e-4 [TV07]. We collect all the binding sites of each PWM $\boldsymbol{P}_j$; if a substring is selected as a binding site by multiple PWMs $\boldsymbol{P}_j$, then we only let the PWM that obtains the highest score use such substring as the binding site. We use the binding sites to re-estimate each PWM $\boldsymbol{P}_j$. Using the newly acquired positions from greedy alignment, we merge these re-estimated PWMs with ALLR and expand/trim using the above procedure.

Although the merged PWMs may closely resemble the real binding sites (see figure 7), we perform greedy alignment instead of directly interpreting the expanded/trimmed PWMs as motifs because the PWMs derived from the triplets may not capture all the binding sites as motifs. We note that this is due to the sparsity induced in convolutional dictionary learning (equation 2) is only homogeneously sparse.

##### B.3 Visualize the filters in the convolutional dictionary learning model

Although the filters learned by uCDL are interpretable, we do not use them directly for motif discovery or indicate them as motifs. Using MA0095.2 from JASPAR as the training set, we show in figure 6 that different batch sizes used during the training of the uCDL network affect the how the filters are learned to represent the input strings. When the batch size is small, the filters are more like k-mers: most columns have high information content. Since k-mers are less expressive, we look at how filters are jointly used to represent substrings in the dataset via the notion of triplets (equation 10). We show in figure 7 that triplets resemble the motifs more clearly than the filters alone. This result matches our intuition that for distributed representation for motifs, we should consider the features *jointly* instead separately (i.e., we should not interpret the features in a distributed representation as if they are local representations).

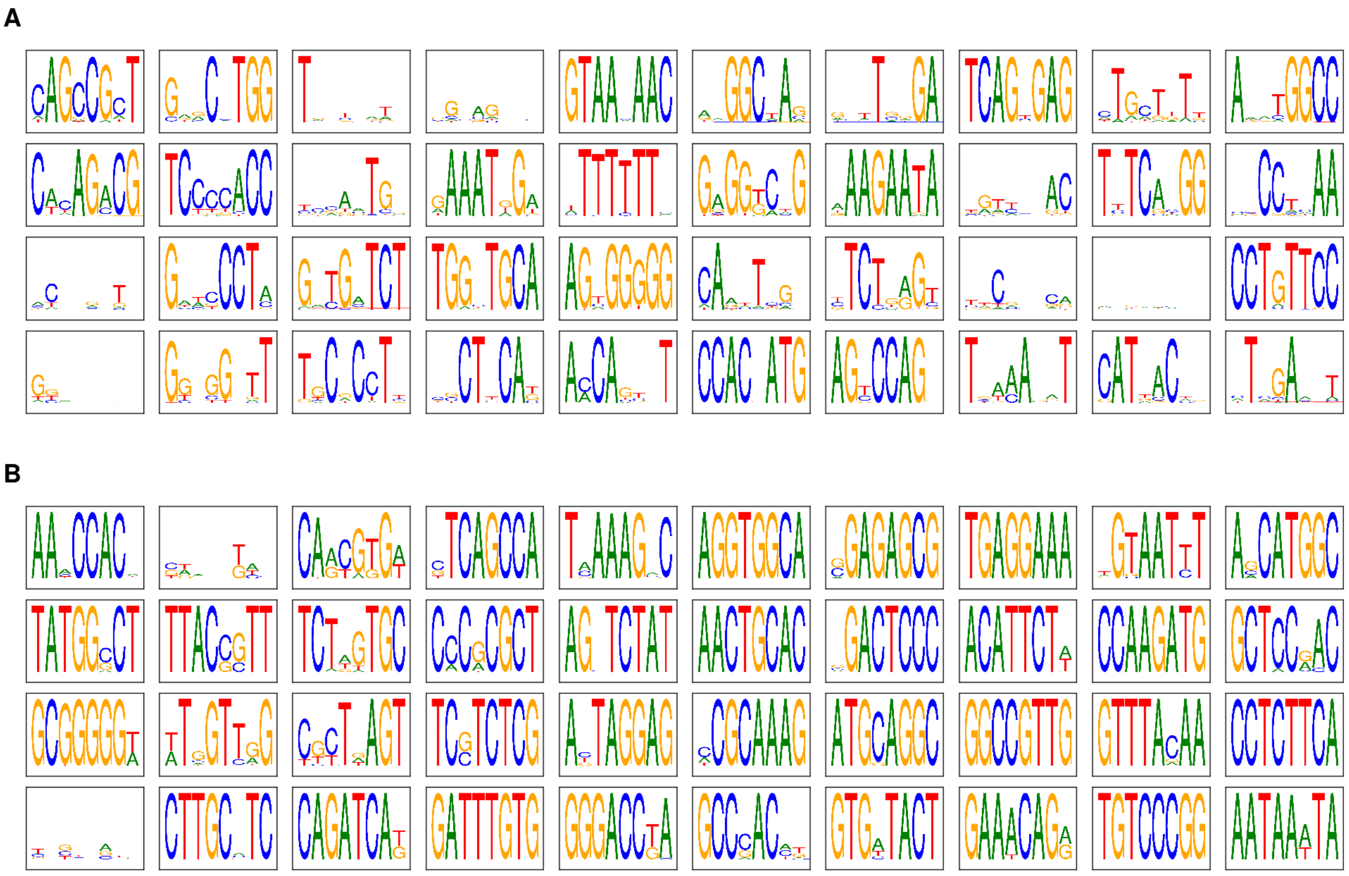

**Figure 6:** Visualizing 40 filters of the dictionary $\boldsymbol{A}$ as PWMs in the unfolded CDL (figure 2) that have high information content, using MA0095.2 from JASPAR as the training set. All the hyperparameters are the same as the setup in B.2 except that we train the network with **A.** batch size set to 200 input strings and **B.** batch size set to 48 input strings.

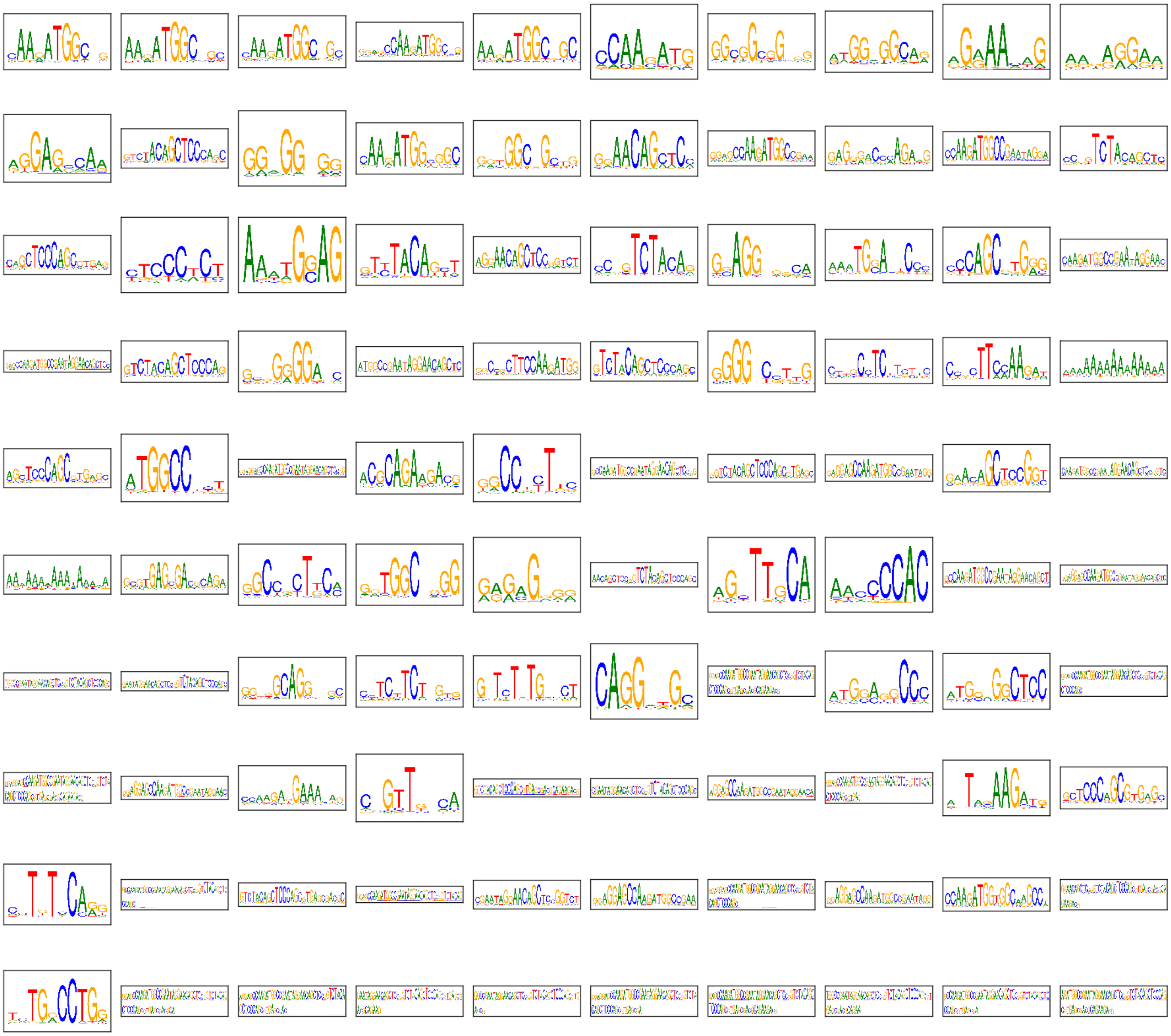

**Figure 7:** Triplets (equation 10), generated from the training result of batch size set to 48 input strings (figure 6-**B**), using MA0095.2 from JASPAR as the training set. The triplets here are the results after expansion using information content followed by merging with average log-likelihood ratio before greedy alignment (section 2.5).

### C Statistical significance of motif enrichment and motifs' co-occurrence

#### C.1 Fisher's exact test for motif enrichment

Following STREME [Bai21], we hold out $10\%$ of the input strings as a test set, left untouched during the training procedure. We evaluate the significance of the motif enrichment on the test set using Fisher's exact test. We create a control set by shuffling each string in the test set that preserves the frequency of 2-mers.

For each $j$-th discovered motif, we do the following: let $N_T = N_t L$ be the number of available positions in the test set, where $N_t$ is the number of strings in the test set and $L$ is the string length. We define the number of "hits" of the $j$-th discovered motif in the test set as $\tau_h$, and the number of "misses" in the test set as $\tau_m = N_T - \tau_h$. Note that a hit at a position is defined by a PWM such that it scores above its score threshold; the score threshold for each PWM using the approximation algorithm pvalue2Score [TV07] with a p-value $2.5\text{e-}4$. We also define the hits and misses for the control set as $c_h$ and $c_m$, respectively. We perform Fisher's exact test of the null hypothesis that the odds ratio $(\tau_h/\tau_m)/(c_h/c_m)$ is one, against the alternative hypothesis that they are not equal.

#### C.2 Pointwise mutual information for motifs' co-occurrence

We estimate the pointwise mutual information (PMI) for each pair of significant motifs and list the pairs whose PMI is greater than zero [Mec+15; II06]. In computational linguistics, the PMI of two words $w_1, w_2$ is defined as

$$\text{PMI}(w_1, w_2) = \log_2 \frac{p(w_1, w_2)}{p(w_1)p(w_2)}$$

where $p(w_1, w_2)$ is the probability of the co-occurrence of two words $w_1$ and $w_2$ and $p(w_1)$, $p(w_2)$ are the probabilities of the occurrence of $w_1$ and $w_2$, respectively. A PMI of a value greater than zero indicates the probability that the co-occurrence of two words is more likely than two words that occur independently.

We estimate the PMI of pairs of significant motifs with the following simple calculations. Suppose we have found $J$ significant motifs. Let $\boldsymbol{C}$ be a $L \times N$ matrix that records whether a motif is present at the $\ell$-th nucleotide of the $n$-th string in the dataset. Specifically, we define each entry $\boldsymbol{c}_{ij}$ of $\boldsymbol{C}$ as

$$\boldsymbol{c}_{\ell n} = \begin{cases} j & \text{if the } j\text{-th motif is present in its forward direction} \\ -j & \text{if the } j\text{-th motif is present in its reverse-complement direction} \end{cases}$$

We define the motif-co-occurrence as

$$f^C(j_1, j_2) = \bar{f}(j_1, j_2) + \bar{f}(-j_2, -j_1), \qquad \text{for } j_1, j_2 \in \{\, 1, 2, \ldots, J, -1, -2, \ldots, -J \,\}$$

where $\bar{f}(j_1, j_2) = |\{\, c_{\ell n} : c_{\ell n} = j_1,\, c_{kn} = j_2,\, c_{\ell n} + l_{j_1} \leq k \leq c_{\ell n} + l_{j_1} + \alpha \,\}|$, $l_{j_1}$ the width of the PWM that represents the $j_1$-th motif, and $\alpha$ is the context width. The motif-co-occurrence $f^C(j_1, j_2)$ shows how many times the $j_2$ motif appears after the $j_1$-th motif within $\alpha$ nucleotides, taking into account both the forward and the reverse complement directions. We thus calculate the PMI for pairs of motifs as follows:

$$\text{PMI}(j_1, j_2) = \log_2 \frac{f^C(j_1, j_2) \sum_{j_1, j_2} f^C(j_1, j_2)}{\sum_{j_1} f^C(j_1, j_2) \sum_{j_2} f^C(j_1, j_2)}$$

for $j_1, j_2$ such that $f^C(j_1, j_2) > 0$. Note that changing the context width $\alpha$ adjust the relationship between pairs of motifs – a lower value of $\alpha$ shows more syntactic relationships, and a higher value of alpha shows more semantic relationships.

For ease of interpretation, we use normalized PMI, defined as

$$\text{nPMI}(j_1, j_2) = \frac{\text{PMI}(j_1, j_2)}{-\log_2(p(j_1, j_2))}.$$

Note that the output value from nPMI falls into the range of $[-1, 1]$ for all $j_1$ and $j_2$. An nPMI of value $-1$ shows that there is no co-occurrence, a value $0$ shows that the pair is independent, and a value of $1$ shows complete co-occurrence.

#### References

[Ros83] Bernard Rosner. "Percentage points for a generalized ESD many-outlier procedure". In: *Technometrics* 25.2 (1983), pp. 165–172.

[SS90] Thomas D Schneider and R Michael Stephens. "Sequence logos: a new way to display consensus sequences". In: *Nucleic acids research* 18.20 (1990), pp. 6097–6100.

[PS+00] Pavel A Pevzner, Sing-Hoi Sze, et al. "Combinatorial approaches to finding subtle signals in DNA sequences." In: *ISMB*. Vol. 8. 2000, pp. 269–278.

[WS03] Ting Wang and Gary D Stormo. "Combining phylogenetic data with co-regulated genes to identify regulatory motifs". In: *Bioinformatics* 19.18 (2003), pp. 2369–2380.

[Cro+04] Gavin E Crooks et al. "WebLogo: a sequence logo generator". In: *Genome research* 14.6 (2004), pp. 1188–1190.

[II06] Aminul Islam and Diana Inkpen. "Second order co-occurrence PMI for determining the semantic similarity of words". In: *Proceedings of the Fifth International Conference on Language Resources and Evaluation (LREC'06)*. 2006.

[TV07] Hélène Touzet and Jean-Stéphane Varré. "Efficient and accurate P-value computation for Position Weight Matrices". In: *Algorithms for Molecular Biology* 2.1 (2007), pp. 1–12.

[Mec+15] Cornelia Meckbach et al. "PC-TraFF: identification of potentially collaborating transcription factors using pointwise mutual information". In: *BMC bioinformatics* 16.1 (2015), pp. 1–21.

[Bez+17] Jeff Bezanson et al. "Julia: A fresh approach to numerical computing". In: *SIAM review* 59.1 (2017), pp. 65–98.

[Woh17] Brendt Wohlberg. "Convolutional sparse coding with overlapping group norms". In: *arXiv preprint arXiv:1708.09038* (2017).

[BFD18] Tim Besard, Christophe Foket, and Bjorn De Sutter. "Effective extensible programming: unleashing Julia on GPUs". In: *IEEE Transactions on Parallel and Distributed Systems* 30.4 (2018), pp. 827–841.

[Inn18] Mike Innes. "Flux: Elegant machine learning with Julia". In: *Journal of Open Source Software* 3.25 (2018), p. 602.

[Bes+19] Mathieu Besançon et al. "Distributions. jl: Definition and modeling of probability distributions in the JuliaStats ecosystem". In: *arXiv preprint arXiv:1907.08611* (2019).

[Zhu+20] Juntang Zhuang et al. "Adabelief optimizer: Adapting stepsizes by the belief in observed gradients". In: *Advances in neural information processing systems* 33 (2020), pp. 18795–18806.

[Bai21] Timothy L Bailey. "STREME: accurate and versatile sequence motif discovery". In: *Bioinformatics* 37.18 (2021), pp. 2834–2840.

[DK21] Simon Danisch and Julius Krumbiegel. "Makie. jl: Flexible high-performance data visualization for Julia". In: *Journal of Open Source Software* 6.65 (2021), p. 3349.
