## Supplemental File 2 for "Deep unfolded convolutional dictionary learning for motif discovery": brief_summary.html

Note that

- Protein name is displayed below each of the Jaspar logo; click the link to go to Jaspar page.
- The statistical significance (p-values from Fisher's exact test) are below each discovered motif.
- This page is just a brief summary. Click the link "More details" on the right for more.

| bzip |  |  |  |  |  |  |  |
| --- | --- | --- | --- | --- | --- | --- | --- |
| Jaspar motif | Data type | Discovered motif 1 | Discovered motif 2 | Discovered motif 3 | Discovered motif 4 | Discovered motif 5 |  |
|  | ChIP-seq |  |  |  |  |  | More details |
| MA0102.3 |  | 3.81e-93 | 8.76e-72 | 1.42e-19 | 1.46e-7 | 0.000935 |  |
|  | ChIP-seq |  |  |  |  |  | More details |
| MA0150.2 |  | 3.23e-19 | 1.17e-7 |  |  |  |  |
|  | ChIP-seq |  |  |  |  |  | More details |
| MA0462.1 |  | 6.73e-29 | 2.67e-11 | 7.78e-5 | 9.37e-5 | 9.42e-5 |  |
|  | ChIP-seq |  |  |  |  |  | More details |
| MA0476.1 |  | 1.61e-81 | 1.11e-41 | 5.38e-30 | 1.45e-22 | 7.54e-14 |  |
|  | ChIP-seq |  |  |  |  |  | More details |
| MA0477.1 |  | 4.16e-96 | 7.96e-10 | 1.04e-9 | 0.00169 |  |  |
|  | ChIP-seq |  |  |  |  |  | More details |
| MA0478.1 |  | 2.91e-48 | 1.48e-35 | 6.56e-14 | 2.89e-11 | 0.00692 |  |
|  | ChIP-seq |  |  |  |  |  | More details |
| MA0488.1 |  | 0.0 | 1.66e-6 | 2.26e-5 |  |  |  |
|  | ChIP-seq |  |  |  |  |  | More details |
| MA0489.1 |  | 9.99e-180 | 3.81e-27 | 1.14e-12 | 4.84e-8 | 4.79e-6 |  |
|  | ChIP-seq |  |  |  |  |  | More details |
| MA0490.1 |  | 7.06e-192 | 2.67e-189 | 2.50e-9 | 6.93e-8 | 0.000277 |  |
|  | ChIP-seq |  |  |  |  |  | More details |
| MA0491.1 |  | 0.0 | 4.48e-96 | 4.55e-6 | 1.14e-5 | 0.000102 |  |
|  | ChIP-seq |  |  |  |  |  | More details |
| MA0492.1 |  | 0.0 | 3.35e-10 | 1.14e-6 | 0.000227 | 0.00256 |  |
|  | ChIP-seq |  |  |  |  |  | More details |
| MA0495.1 |  | 0.0 | 1.26e-272 | 2.34e-19 | 3.21e-8 | 1.48e-5 |  |
|  | ChIP-seq |  |  |  |  |  | More details |
| MA0496.1 |  | 0.0 | 0.0 | 1.95e-5 | 3.51e-5 | 8.85e-5 |  |
|  | ChIP-seq |  |  |  |  |  | More details |
| MA0501.1 |  | 1.68e-33 |  |  |  |  |  |
|  | ChIP-seq |  |  |  |  |  | More details |
| MA0506.1 |  | 3.02e-66 | 0.00433 |  |  |  |  |
|  | ChIP-chip |  |  |  |  |  | More details |
| MA0530.1 |  |  |  |  |  |  |  |
|  | ChIP-seq |  |  |  |  |  | More details |
| MA0547.1 |  | 3.22e-5 | 0.00373 |  |  |  |  |
|  | SELEX |  |  |  |  |  | More details |
| MA0588.1 |  | 1.80e-13 |  |  |  |  |  |
|  | ChIP-seq |  |  |  |  |  | More details |
| MA0591.1 |  | 4.38e-7 |  |  |  |  |  |
|  | PBM |  |  |  |  |  | More details |
| MA0931.1 |  | 7.44e-9 | 3.37e-6 |  |  |  |  |
|  | PBM |  |  |  |  |  | More details |
| MA0941.1 |  | 5.22e-12 |  |  |  |  |  |
|  | PBM |  |  |  |  |  | More details |
| MA1047.1 |  | 8.28e-15 | 0.008 |  |  |  |  |
|  | PBM |  |  |  |  |  | More details |
| MA1068.1 |  | 5.53e-17 |  |  |  |  |  |
|  | PBM |  |  |  |  |  | More details |
| MA1069.1 |  | 9.71e-13 |  |  |  |  |  |
|  | DAP-seq |  |  |  |  |  | More details |
| MA1334.1 |  | 6.05e-18 |  |  |  |  |  |
|  | DAP-seq |  |  |  |  |  | More details |
| MA1335.1 |  | 1.58e-7 | 1.97e-5 |  |  |  |  |
|  | DAP-seq |  |  |  |  |  | More details |
| MA1336.1 |  | 4.18e-14 |  |  |  |  |  |
|  | DAP-seq |  |  |  |  |  | More details |
| MA1337.1 |  | 1.06e-13 |  |  |  |  |  |
|  | DAP-seq |  |  |  |  |  | More details |
| MA1338.1 |  | 5.07e-20 |  |  |  |  |  |
|  | DAP-seq |  |  |  |  |  | More details |
| MA1340.1 |  | 7.25e-16 |  |  |  |  |  |
|  | DAP-seq |  |  |  |  |  | More details |
| MA1341.1 |  | 9.89e-17 |  |  |  |  |  |
|  | DAP-seq |  |  |  |  |  | More details |
| MA1343.1 |  |  |  |  |  |  |  |
|  | DAP-seq |  |  |  |  |  | More details |
| MA1345.1 |  | 3.47e-16 |  |  |  |  |  |
|  | DAP-seq |  |  |  |  |  | More details |
| MA1346.1 |  | 2.04e-7 | 7.51e-6 | 0.00131 |  |  |  |
|  | DAP-seq |  |  |  |  |  | More details |
| MA1348.1 |  | 1.84e-12 |  |  |  |  |  |
|  | DAP-seq |  |  |  |  |  | More details |
| MA1349.1 |  |  |  |  |  |  |  |
|  | ChIP-seq |  |  |  |  |  | More details |
| MA1351.1 |  | 2.34e-16 |  |  |  |  |  |

| stat |  |  |  |  |  |  |  |
| --- | --- | --- | --- | --- | --- | --- | --- |
| Jaspar motif | Data type | Discovered motif 1 | Discovered motif 2 | Discovered motif 3 | Discovered motif 4 | Discovered motif 5 |  |
|  | ChIP-seq |  |  |  |  |  | More details |
| MA0137.3 |  | 1.69e-26 | 3.76e-25 | 1.36e-17 | 7.40e-10 | 2.02e-9 |  |
|  | ChIP-seq |  |  |  |  |  | More details |
| MA0144.2 |  | 2.57e-197 | 5.16e-32 | 7.80e-20 | 3.88e-16 | 1.90e-12 |  |
|  | ChIP-seq |  |  |  |  |  | More details |
| MA0517.1 |  | 1.47e-14 |  |  |  |  |  |
|  | ChIP-seq |  |  |  |  |  | More details |
| MA0518.1 |  | 1.66e-61 | 0.000146 | 0.000267 | 0.00957 |  |  |
|  | ChIP-seq |  |  |  |  |  | More details |
| MA0519.1 |  | 2.42e-316 | 3.95e-5 | 0.000376 | 0.000899 | 0.00352 |  |
|  | ChIP-seq |  |  |  |  |  | More details |
| MA0520.1 |  | 2.99e-40 | 7.15e-7 | 0.00546 | 0.0059 |  |  |
|  | ChIP-chip |  |  |  |  |  | More details |
| MA0532.1 |  |  |  |  |  |  |  |

| forkhead\_helix\_factor |  |  |  |  |  |  |  |
| --- | --- | --- | --- | --- | --- | --- | --- |
| Jaspar motif | Data type | Discovered motif 1 | Discovered motif 2 | Discovered motif 3 | Discovered motif 4 | Discovered motif 5 |  |
|  | ChIP-seq |  |  |  |  |  | More details |
| MA0024.2 |  | 0.00252 |  |  |  |  |  |
|  | ChIP-seq |  |  |  |  |  | More details |
| MA0148.3 |  | 0.0 | 1.26e-35 | 1.69e-7 | 5.44e-6 | 0.00113 |  |
|  | ChIP-seq |  |  |  |  |  | More details |
| MA0470.1 |  | 3.43e-21 | 9.82e-5 |  |  |  |  |
|  | ChIP-seq |  |  |  |  |  | More details |
| MA0471.1 |  | 1.92e-17 | 6.17e-7 | 0.00528 |  |  |  |
|  | ChIP-seq |  |  |  |  |  | More details |
| MA0479.1 |  | 3.30e-105 | 1.72e-10 | 0.00208 |  |  |  |
|  | ChIP-seq |  |  |  |  |  | More details |
| MA0480.1 |  | 1.06e-18 | 3.65e-17 | 6.04e-7 | 5.57e-5 |  |  |
|  | ChIP-seq |  |  |  |  |  | More details |
| MA0481.1 |  | 7.65e-7 |  |  |  |  |  |
|  | ChIP-seq |  |  |  |  |  | More details |
| MA0509.1 |  | 1.94e-42 | 3.59e-21 | 0.000228 |  |  |  |
|  | ChIP-seq |  |  |  |  |  | More details |
| MA0510.1 |  | 3.21e-32 | 7.98e-23 | 3.33e-12 | 0.00113 |  |  |
|  | ChIP-seq |  |  |  |  |  | More details |
| MA0541.1 |  | 2.59e-17 | 1.71e-16 | 8.23e-8 | 0.000302 |  |  |
|  | ChIP-seq |  |  |  |  |  | More details |
| MA0546.1 |  | 2.33e-8 | 2.63e-6 |  |  |  |  |
|  | ChIP-seq |  |  |  |  |  | More details |
| MA0593.1 |  | 9.32e-13 | 3.74e-7 |  |  |  |  |
|  | ChIP-seq |  |  |  |  |  | More details |
| MA0600.1 |  | 1.40e-54 | 3.41e-13 | 0.000122 | 0.00224 |  |  |

| other\_C4\_ZF |  |  |  |  |  |  |  |
| --- | --- | --- | --- | --- | --- | --- | --- |
| Jaspar motif | Data type | Discovered motif 1 | Discovered motif 2 | Discovered motif 3 | Discovered motif 4 | Discovered motif 5 |  |
|  | ChIP-seq |  |  |  |  |  | More details |
| MA0036.2 |  | 3.38e-33 | 3.48e-12 | 4.87e-11 | 0.00269 | 0.0031 |  |
|  | ChIP-seq |  |  |  |  |  | More details |
| MA0140.2 |  | 6.61e-90 | 1.36e-17 | 5.85e-10 |  |  |  |
|  | ChIP-seq |  |  |  |  |  | More details |
| MA0482.1 |  | 3.86e-41 | 5.77e-7 | 3.85e-5 |  |  |  |
|  | ChIP-chip |  |  |  |  |  | More details |
| MA0536.1 |  | 1.30e-20 | 0.00395 |  |  |  |  |
|  | ChIP-seq |  |  |  |  |  | More details |
| MA0538.1 |  | 0.000964 |  |  |  |  |  |
|  | PBM |  |  |  |  |  | More details |
| MA0974.1 |  |  |  |  |  |  |  |
|  | PBM |  |  |  |  |  | More details |
| MA0981.1 |  | 0.00112 |  |  |  |  |  |
|  | PBM |  |  |  |  |  | More details |
| MA0982.1 |  | 0.00106 | 0.00919 |  |  |  |  |
|  | PBM |  |  |  |  |  | More details |
| MA0983.1 |  | 0.00183 |  |  |  |  |  |
|  | PBM |  |  |  |  |  | More details |
| MA1014.1 |  | 1.24e-13 |  |  |  |  |  |
|  | DAP-seq |  |  |  |  |  | More details |
| MA1267.1 |  | 0.00319 |  |  |  |  |  |
|  | DAP-seq |  |  |  |  |  | More details |
| MA1268.1 |  | 0.0051 |  |  |  |  |  |
|  | DAP-seq |  |  |  |  |  | More details |
| MA1269.1 |  | 0.00387 |  |  |  |  |  |
|  | DAP-seq |  |  |  |  |  | More details |
| MA1270.1 |  | 1.20e-5 |  |  |  |  |  |
|  | DAP-seq |  |  |  |  |  | More details |
| MA1272.1 |  | 0.00186 |  |  |  |  |  |
|  | DAP-seq |  |  |  |  |  | More details |
| MA1273.1 |  | 0.0001 |  |  |  |  |  |
|  | DAP-seq |  |  |  |  |  | More details |
| MA1274.1 |  |  |  |  |  |  |  |
|  | DAP-seq |  |  |  |  |  | More details |
| MA1275.1 |  | 2.07e-6 |  |  |  |  |  |
|  | DAP-seq |  |  |  |  |  | More details |
| MA1276.1 |  | 2.66e-6 | 0.00107 |  |  |  |  |
|  | DAP-seq |  |  |  |  |  | More details |
| MA1277.1 |  | 4.32e-5 |  |  |  |  |  |
|  | DAP-seq |  |  |  |  |  | More details |
| MA1278.1 |  |  |  |  |  |  |  |
|  | DAP-seq |  |  |  |  |  | More details |
| MA1279.1 |  | 7.47e-5 |  |  |  |  |  |
|  | DAP-seq |  |  |  |  |  | More details |
| MA1280.1 |  |  |  |  |  |  |  |
|  | DAP-seq |  |  |  |  |  | More details |
| MA1281.1 |  |  |  |  |  |  |  |
|  | DAP-seq |  |  |  |  |  | More details |
| MA1323.1 |  | 0.000196 |  |  |  |  |  |

| MADS\_box\_factors |  |  |  |  |  |  |  |
| --- | --- | --- | --- | --- | --- | --- | --- |
| Jaspar motif | Data type | Discovered motif 1 | Discovered motif 2 | Discovered motif 3 | Discovered motif 4 | Discovered motif 5 |  |
|  | ChIP-seq |  |  |  |  |  | More details |
| MA0052.2 |  | 1.98e-35 | 0.00221 | 0.00614 |  |  |  |
|  | ChIP-seq |  |  |  |  |  | More details |
| MA0083.2 |  | 2.64e-30 | 6.44e-19 |  |  |  |  |
|  | ChIP-seq |  |  |  |  |  | More details |
| MA0497.1 |  | 2.09e-24 | 1.67e-12 | 0.000112 | 0.00191 | 0.00987 |  |
|  | ChIP-chip |  |  |  |  |  | More details |
| MA0548.1 |  |  |  |  |  |  |  |
|  | DAP-seq |  |  |  |  |  | More details |
| MA0548.2 |  | 3.17e-6 |  |  |  |  |  |
|  | ChIP-chip |  |  |  |  |  | More details |
| MA0554.1 |  | 1.94e-8 |  |  |  |  |  |
|  | ChIP-seq |  |  |  |  |  | More details |
| MA0556.1 |  | 0.000427 |  |  |  |  |  |
|  | ChIP-seq |  |  |  |  |  | More details |
| MA0558.1 |  |  |  |  |  |  |  |
|  | ChIP-seq |  |  |  |  |  | More details |
| MA0559.1 |  | 0.000713 | 0.00296 |  |  |  |  |
|  | ChIP-seq |  |  |  |  |  | More details |
| MA0563.1 |  |  |  |  |  |  |  |
|  | ChIP-seq |  |  |  |  |  | More details |
| MA0940.1 |  | 6.98e-7 | 0.00292 | 0.00496 |  |  |  |
|  | ChIP-seq |  |  |  |  |  | More details |
| MA1012.1 |  |  |  |  |  |  |  |
|  | DAP-seq |  |  |  |  |  | More details |
| MA1203.1 |  | 7.11e-16 |  |  |  |  |  |

| helix\_span\_helix |  |  |  |  |  |  |  |
| --- | --- | --- | --- | --- | --- | --- | --- |
| Jaspar motif | Data type | Discovered motif 1 | Discovered motif 2 | Discovered motif 3 | Discovered motif 4 | Discovered motif 5 |  |
|  | ChIP-seq |  |  |  |  |  | More details |
| MA0003.2 |  | 1.24e-34 | 9.51e-5 | 0.00115 | 0.00273 | 0.0085 |  |
|  | ChIP-seq |  |  |  |  |  | More details |
| MA0104.3 |  | 2.86e-20 |  |  |  |  |  |

| tryptophan\_cluster\_factors |  |  |  |  |  |  |  |
| --- | --- | --- | --- | --- | --- | --- | --- |
| Jaspar motif | Data type | Discovered motif 1 | Discovered motif 2 | Discovered motif 3 | Discovered motif 4 | Discovered motif 5 |  |
|  | ChIP-seq |  |  |  |  |  | More details |
| MA0050.2 |  | 1.17e-35 |  |  |  |  |  |
|  | ChIP-seq |  |  |  |  |  | More details |
| MA0076.2 |  | 2.32e-47 | 6.62e-6 |  |  |  |  |
|  | ChIP-seq |  |  |  |  |  | More details |
| MA0098.2 |  | 7.96e-33 | 1.59e-10 | 1.32e-7 |  |  |  |
|  | ChIP-seq |  |  |  |  |  | More details |
| MA0100.2 |  | 9.99e-14 | 1.47e-6 |  |  |  |  |
|  | ChIP-seq |  |  |  |  |  | More details |
| MA0473.1 |  | 3.84e-103 | 4.41e-57 | 4.34e-18 | 3.91e-17 | 1.86e-16 |  |
|  | ChIP-seq |  |  |  |  |  | More details |
| MA0475.1 |  | 1.79e-24 | 1.68e-22 | 2.59e-20 | 7.38e-5 | 0.00944 |  |
|  | ChIP-seq |  |  |  |  |  | More details |
| MA0544.1 |  | 4.34e-10 | 0.00779 |  |  |  |  |
|  | ChIP-seq |  |  |  |  |  | More details |
| MA0598.1 |  | 1.53e-10 | 5.73e-6 | 0.000773 |  |  |  |
|  | PBM |  |  |  |  |  | More details |
| MA1020.1 |  | 9.57e-13 |  |  |  |  |  |
|  | DAP-seq |  |  |  |  |  | More details |
| MA1163.1 |  | 7.48e-12 |  |  |  |  |  |
|  | DAP-seq |  |  |  |  |  | More details |
| MA1164.1 |  | 9.50e-6 | 2.91e-5 |  |  |  |  |
|  | DAP-seq |  |  |  |  |  | More details |
| MA1165.1 |  | 3.55e-10 | 2.13e-7 | 0.00223 |  |  |  |
|  | DAP-seq |  |  |  |  |  | More details |
| MA1167.1 |  | 2.54e-7 | 4.20e-5 |  |  |  |  |
|  | DAP-seq |  |  |  |  |  | More details |
| MA1168.1 |  | 9.03e-10 |  |  |  |  |  |
|  | DAP-seq |  |  |  |  |  | More details |
| MA1172.1 |  |  |  |  |  |  |  |
|  | DAP-seq |  |  |  |  |  | More details |
| MA1173.1 |  | 1.51e-6 | 0.000707 |  |  |  |  |
|  | DAP-seq |  |  |  |  |  | More details |
| MA1174.1 |  | 3.23e-14 |  |  |  |  |  |
|  | DAP-seq |  |  |  |  |  | More details |
| MA1175.1 |  | 6.52e-7 |  |  |  |  |  |
|  | DAP-seq |  |  |  |  |  | More details |
| MA1176.1 |  | 6.17e-14 |  |  |  |  |  |
|  | DAP-seq |  |  |  |  |  | More details |
| MA1177.1 |  | 3.75e-9 |  |  |  |  |  |
|  | DAP-seq |  |  |  |  |  | More details |
| MA1178.1 |  |  |  |  |  |  |  |
|  | DAP-seq |  |  |  |  |  | More details |
| MA1180.1 |  |  |  |  |  |  |  |
|  | DAP-seq |  |  |  |  |  | More details |
| MA1182.1 |  | 8.17e-11 | 2.12e-6 |  |  |  |  |
|  | DAP-seq |  |  |  |  |  | More details |
| MA1183.1 |  | 0.00019 |  |  |  |  |  |
|  | DAP-seq |  |  |  |  |  | More details |
| MA1184.1 |  | 2.05e-9 | 3.19e-9 |  |  |  |  |
|  | DAP-seq |  |  |  |  |  | More details |
| MA1185.1 |  | 8.59e-9 |  |  |  |  |  |
|  | DAP-seq |  |  |  |  |  | More details |
| MA1186.1 |  | 3.26e-11 | 2.01e-5 | 0.000191 | 0.000707 |  |  |
|  | DAP-seq |  |  |  |  |  | More details |
| MA1187.1 |  | 1.94e-10 |  |  |  |  |  |
|  | DAP-seq |  |  |  |  |  | More details |
| MA1188.1 |  | 0.000211 | 0.000433 | 0.00305 |  |  |  |
|  | DAP-seq |  |  |  |  |  | More details |
| MA1189.1 |  | 8.08e-9 | 1.73e-7 |  |  |  |  |
|  | DAP-seq |  |  |  |  |  | More details |
| MA1190.1 |  | 6.64e-9 | 2.81e-5 |  |  |  |  |
|  | DAP-seq |  |  |  |  |  | More details |
| MA1191.1 |  | 4.57e-7 |  |  |  |  |  |
|  | DAP-seq |  |  |  |  |  | More details |
| MA1195.1 |  | 1.15e-10 |  |  |  |  |  |
|  | DAP-seq |  |  |  |  |  | More details |
| MA1196.1 |  | 3.29e-7 | 0.000707 |  |  |  |  |
|  | DAP-seq |  |  |  |  |  | More details |
| MA1207.1 |  |  |  |  |  |  |  |
|  | DAP-seq |  |  |  |  |  | More details |
| MA1292.1 |  | 1.46e-15 |  |  |  |  |  |
|  | DAP-seq |  |  |  |  |  | More details |
| MA1293.1 |  | 0.00758 |  |  |  |  |  |
|  | DAP-seq |  |  |  |  |  | More details |
| MA1352.1 |  | 2.51e-6 |  |  |  |  |  |
|  | DAP-seq |  |  |  |  |  | More details |
| MA1353.1 |  | 3.57e-12 | 1.36e-6 |  |  |  |  |
|  | DAP-seq |  |  |  |  |  | More details |
| MA1355.1 |  | 3.42e-14 | 2.92e-7 |  |  |  |  |
|  | DAP-seq |  |  |  |  |  | More details |
| MA1356.1 |  | 5.99e-13 |  |  |  |  |  |
|  | DAP-seq |  |  |  |  |  | More details |
| MA1366.1 |  | 6.63e-6 | 0.000144 |  |  |  |  |
|  | DAP-seq |  |  |  |  |  | More details |
| MA1384.1 |  | 5.90e-11 |  |  |  |  |  |
|  | DAP-seq |  |  |  |  |  | More details |
| MA1385.1 |  | 1.39e-8 |  |  |  |  |  |
|  | DAP-seq |  |  |  |  |  | More details |
| MA1386.1 |  | 8.31e-20 |  |  |  |  |  |
|  | DAP-seq |  |  |  |  |  | More details |
| MA1388.1 |  | 6.06e-9 | 0.000153 | 0.00216 | 0.00316 |  |  |
|  | DAP-seq |  |  |  |  |  | More details |
| MA1389.1 |  | 2.47e-15 |  |  |  |  |  |
|  | DAP-seq |  |  |  |  |  | More details |
| MA1390.1 |  | 1.66e-16 |  |  |  |  |  |
|  | DAP-seq |  |  |  |  |  | More details |
| MA1391.1 |  | 2.62e-11 | 3.50e-5 |  |  |  |  |
|  | DAP-seq |  |  |  |  |  | More details |
| MA1392.1 |  | 9.21e-9 |  |  |  |  |  |
|  | DAP-seq |  |  |  |  |  | More details |
| MA1393.1 |  | 8.36e-12 |  |  |  |  |  |
|  | DAP-seq |  |  |  |  |  | More details |
| MA1394.1 |  | 5.28e-10 | 2.56e-6 | 0.00528 |  |  |  |
|  | DAP-seq |  |  |  |  |  | More details |
| MA1397.1 |  | 1.77e-16 | 2.26e-5 |  |  |  |  |
|  | DAP-seq |  |  |  |  |  | More details |
| MA1398.1 |  | 4.25e-7 | 0.000139 |  |  |  |  |
|  | DAP-seq |  |  |  |  |  | More details |
| MA1400.1 |  | 2.64e-7 |  |  |  |  |  |
|  | DAP-seq |  |  |  |  |  | More details |
| MA1401.1 |  | 1.21e-9 |  |  |  |  |  |

| paired\_box\_factors |  |  |  |  |  |  |  |
| --- | --- | --- | --- | --- | --- | --- | --- |
| Jaspar motif | Data type | Discovered motif 1 | Discovered motif 2 | Discovered motif 3 | Discovered motif 4 | Discovered motif 5 |  |
|  | ChIP-seq |  |  |  |  |  | More details |
| MA0014.2 |  | 0.000258 | 0.0039 |  |  |  |  |

| p53\_domain\_factors |  |  |  |  |  |  |  |
| --- | --- | --- | --- | --- | --- | --- | --- |
| Jaspar motif | Data type | Discovered motif 1 | Discovered motif 2 | Discovered motif 3 | Discovered motif 4 | Discovered motif 5 |  |
|  | ChIP-seq |  |  |  |  |  | More details |
| MA0106.2 |  | 5.01e-25 | 1.16e-10 |  |  |  |  |
|  | ChIP-seq |  |  |  |  |  | More details |
| MA0525.1 |  | 7.67e-88 | 2.47e-22 | 2.14e-20 | 1.57e-12 | 3.25e-6 |  |
|  | DAP-seq |  |  |  |  |  | More details |
| MA1341.1 |  | 1.35e-10 | 7.25e-9 | 0.00375 |  |  |  |

| rel\_homology\_region |  |  |  |  |  |  |  |
| --- | --- | --- | --- | --- | --- | --- | --- |
| Jaspar motif | Data type | Discovered motif 1 | Discovered motif 2 | Discovered motif 3 | Discovered motif 4 | Discovered motif 5 |  |
|  | ChIP-seq |  |  |  |  |  | More details |
| MA0105.3 |  | 7.71e-92 | 0.000121 |  |  |  |  |
|  | ChIP-seq |  |  |  |  |  | More details |
| MA0154.2 |  | 3.56e-237 | 6.10e-220 | 2.62e-14 | 2.88e-10 | 3.24e-8 |  |

| c2h2 |  |  |  |  |  |  |  |
| --- | --- | --- | --- | --- | --- | --- | --- |
| Jaspar motif | Data type | Discovered motif 1 | Discovered motif 2 | Discovered motif 3 | Discovered motif 4 | Discovered motif 5 |  |
|  | ChIP-seq |  |  |  |  |  | More details |
| MA0079.3 |  | 6.03e-48 | 2.05e-18 | 4.33e-10 |  |  |  |
|  | ChIP-seq |  |  |  |  |  | More details |
| MA0095.2 |  | 1.67e-102 | 1.08e-16 | 1.75e-11 | 1.90e-11 | 2.08e-7 |  |
|  | ChIP-seq |  |  |  |  |  | More details |
| MA0162.2 |  | 1.95e-40 | 9.11e-22 | 2.76e-19 | 2.11e-7 | 1.53e-6 |  |
|  | ChIP-chip |  |  |  |  |  | More details |
| MA0452.2 |  | 1.47e-21 |  |  |  |  |  |
|  | ChIP-seq |  |  |  |  |  | More details |
| MA0463.1 |  | 4.88e-12 | 0.00254 |  |  |  |  |
|  | ChIP-seq |  |  |  |  |  | More details |
| MA0472.1 |  | 1.82e-6 | 0.00674 |  |  |  |  |
|  | ChIP-seq |  |  |  |  |  | More details |
| MA0483.1 |  | 2.72e-18 | 3.23e-16 |  |  |  |  |
|  | ChIP-seq |  |  |  |  |  | More details |
| MA0493.1 |  | 1.05e-8 | 0.000263 |  |  |  |  |
|  | ChIP-seq |  |  |  |  |  | More details |
| MA0508.1 |  | 6.37e-52 | 1.54e-12 | 8.11e-7 | 0.000596 | 0.0018 |  |
|  | ChIP-seq |  |  |  |  |  | More details |
| MA0516.1 |  | 3.98e-25 | 1.04e-5 | 1.12e-5 | 0.000253 | 0.00416 |  |
|  | ChIP-seq |  |  |  |  |  | More details |
| MA0527.1 |  | 8.03e-7 |  |  |  |  |  |
|  | ChIP-seq |  |  |  |  |  | More details |
| MA0528.1 |  | 4.11e-15 | 0.000144 | 0.00107 |  |  |  |
|  | ChIP-chip |  |  |  |  |  | More details |
| MA0529.1 |  | 8.74e-52 | 0.000502 |  |  |  |  |
|  | ChIP-chip |  |  |  |  |  | More details |
| MA0531.1 |  | 3.54e-27 | 6.70e-14 |  |  |  |  |
|  | ChIP-chip |  |  |  |  |  | More details |
| MA0533.1 |  | 3.04e-55 | 1.60e-32 | 4.77e-12 | 5.10e-8 | 0.00104 |  |
|  | ChIP-seq |  |  |  |  |  | More details |
| MA0537.1 |  | 3.92e-8 | 1.34e-7 | 3.49e-7 | 0.000818 |  |  |
|  | ChIP-seq |  |  |  |  |  | More details |
| MA0543.1 |  | 2.36e-18 |  |  |  |  |  |
|  | DAP-seq |  |  |  |  |  | More details |
| MA0586.2 |  | 7.38e-18 |  |  |  |  |  |
|  | ChIP-seq |  |  |  |  |  | More details |
| MA0599.1 |  | 4.44e-69 | 5.93e-33 | 9.76e-30 | 0.000323 |  |  |
|  | PBM |  |  |  |  |  | More details |
| MA1055.1 |  |  |  |  |  |  |  |
|  | DAP-seq |  |  |  |  |  | More details |
| MA1059.2 |  | 1.66e-17 |  |  |  |  |  |
|  | DAP-seq |  |  |  |  |  | More details |
| MA1156.1 |  | 6.13e-12 |  |  |  |  |  |
|  | DAP-seq |  |  |  |  |  | More details |
| MA1159.1 |  | 1.02e-13 |  |  |  |  |  |
|  | DAP-seq |  |  |  |  |  | More details |
| MA1321.1 |  | 5.56e-11 | 2.75e-5 | 0.000263 | 0.0027 |  |  |
|  | DAP-seq |  |  |  |  |  | More details |
| MA1322.1 |  | 4.49e-16 | 0.000424 |  |  |  |  |
|  | DAP-seq |  |  |  |  |  | More details |
| MA1371.1 |  | 1.38e-9 |  |  |  |  |  |
|  | DAP-seq |  |  |  |  |  | More details |
| MA1372.1 |  | 1.19e-7 | 0.00453 |  |  |  |  |
|  | DAP-seq |  |  |  |  |  | More details |
| MA1374.1 |  | 3.15e-10 |  |  |  |  |  |

| nuclear\_receptors\_w\_C4 |  |  |  |  |  |  |  |
| --- | --- | --- | --- | --- | --- | --- | --- |
| Jaspar motif | Data type | Discovered motif 1 | Discovered motif 2 | Discovered motif 3 | Discovered motif 4 | Discovered motif 5 |  |
|  | ChIP-seq |  |  |  |  |  | More details |
| MA0007.2 |  | 5.03e-109 | 5.17e-95 | 2.11e-20 | 1.05e-7 | 1.22e-7 |  |
|  | ChIP-seq |  |  |  |  |  | More details |
| MA0113.2 |  | 1.18e-7 |  |  |  |  |  |
|  | ChIP-seq |  |  |  |  |  | More details |
| MA0114.2 |  | 0.0 | 4.90e-6 | 0.000211 | 0.00608 |  |  |
|  | ChIP-seq |  |  |  |  |  | More details |
| MA0258.2 |  | 1.27e-64 | 4.71e-41 | 1.22e-34 | 1.73e-5 |  |  |
|  | ChIP-seq |  |  |  |  |  | More details |
| MA0484.1 |  | 8.17e-86 | 1.06e-77 | 2.86e-25 | 0.000496 |  |  |
|  | ChIP-seq |  |  |  |  |  | More details |
| MA0494.1 |  | 1.00e-24 | 0.00216 |  |  |  |  |
|  | ChIP-seq |  |  |  |  |  | More details |
| MA0504.1 |  | 1.89e-10 |  |  |  |  |  |
|  | ChIP-seq |  |  |  |  |  | More details |
| MA0505.1 |  | 1.17e-37 | 1.12e-7 |  |  |  |  |
|  | ChIP-seq |  |  |  |  |  | More details |
| MA0512.1 |  | 9.75e-65 | 6.35e-25 | 9.39e-10 | 1.24e-7 |  |  |
|  | ChIP-seq |  |  |  |  |  | More details |
| MA0534.1 |  |  |  |  |  |  |  |
|  | ChIP-seq |  |  |  |  |  | More details |
| MA0592.1 |  | 4.74e-7 |  |  |  |  |  |
