## Supplemental File 2 for "Deep unfolded convolutional dictionary learning for motif discovery": weblogo render.ipynb

```
# %% [code]
import os
mat = [i for i in os.listdir() if 'transfac' in i]

# %% [code]
mat_name = [i.split(".t")[0] for i in mat]

# %% [code]
mat

# %% [code]
for ind, m in enumerate(mat):
    cmd = "weblogo -D transfac -f " +m+" --errorbars NO -F png --fineprint \" \" --resolution 60 -s large --fontsize 17 --color-scheme classic  -o " +mat_name[ind]+".png"
    os.system(cmd)

# %% [code]
# weblogo -f MA0079.3.transfac -F png --fineprint " " --resolution 600 -s large --fontsize 17 --color-scheme classic -Y NO -o d1.png
```
