## Supplemental File 2 for "Deep unfolded convolutional dictionary learning for motif discovery": weblogo_render.py

```
import os
mat = [i for i in os.listdir() if 'transfac' in i]
mat_name = [i.split(".t")[0] for i in mat]
for ind, m in enumerate(mat):
    cmd = "weblogo -D transfac -f " +m+" --errorbars NO -F png --fineprint \" \" --resolution 60 -s large --fontsize 17 --color-scheme classic  -o " +mat_name[ind]+".png"
    os.system(cmd)
```
