## Supplementary figures and images for "Deep unfolded convolutional dictionary learning for motif discovery"

### MA0003.2.png

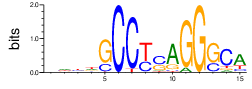

### MA0007.2.png

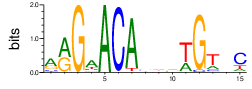

### MA0014.2.png

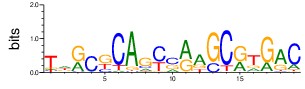

### MA0024.2.png

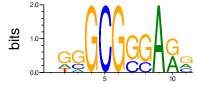

### MA0052.2.png

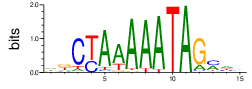

### MA0058.2.png

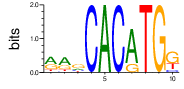

### MA0083.2.png

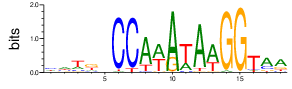

### MA0093.2.png

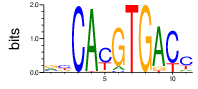

### MA0102.3.png

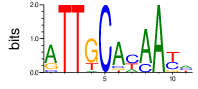

### MA0104.3.png

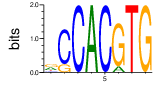

### MA0105.3.png

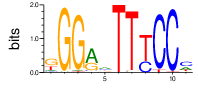

### MA0106.2.png

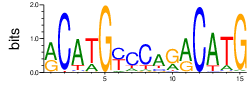

### MA0113.2.png

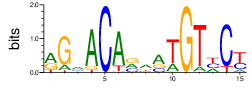

### MA0114.2.png

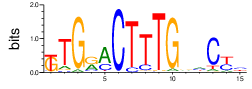

### MA0137.3.png

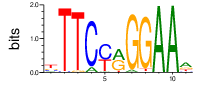

### MA0144.2.png

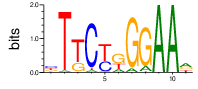

### MA0148.3.png

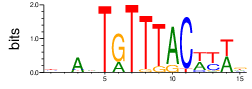

### MA0150.2.png

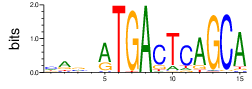

### MA0154.2.png

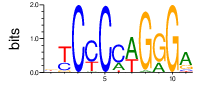

### MA0258.2.png

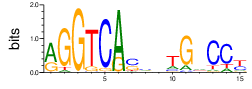

### MA0462.1.png

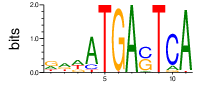

### MA0466.1.png

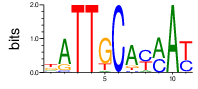

### MA0470.1.png

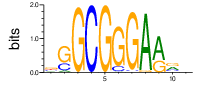

### MA0471.1.png

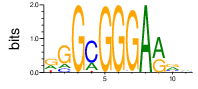

### MA0476.1.png

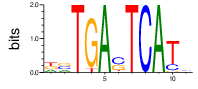

### MA0477.1.png

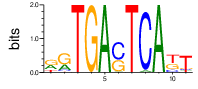

### MA0478.1.png

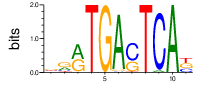

### MA0479.1.png

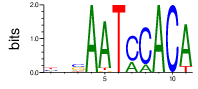

### MA0480.1.png

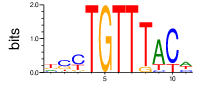

### MA0481.1.png

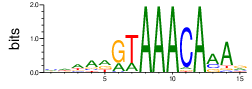

### MA0484.1.png

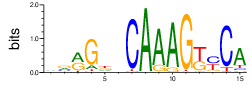

### MA0488.1.png

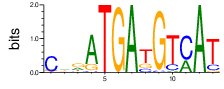

### MA0489.1.png

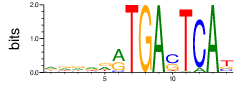

### MA0490.1.png

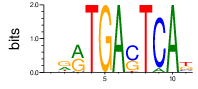

### MA0491.1.png

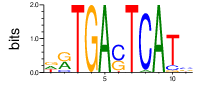

### MA0492.1.png

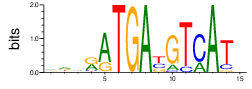

### MA0494.1.png

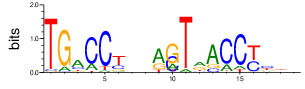

### MA0495.1.png

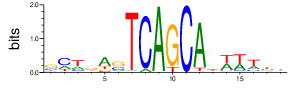

### MA0496.1.png

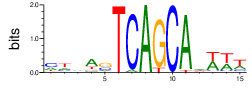

### MA0497.1.png

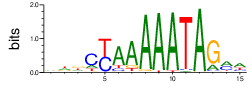

### MA0501.1.png

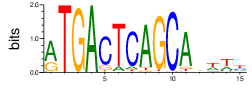

### MA0504.1.png

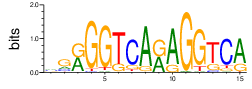

### MA0505.1.png

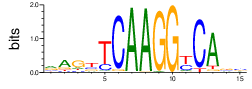

### MA0506.1.png

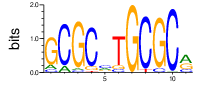

### MA0509.1.png

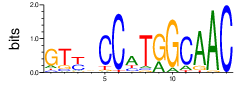

### MA0510.1.png

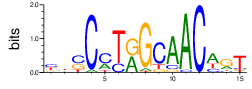

### MA0512.1.png

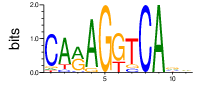

### MA0517.1.png

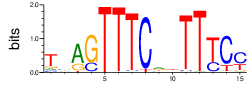

### MA0518.1.png

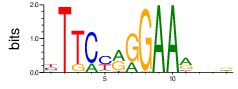

### MA0519.1.png

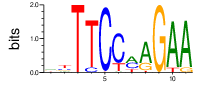

### MA0520.1.png

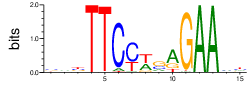

### MA0524.1.png

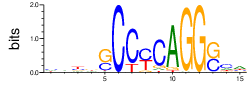

### MA0525.1.png

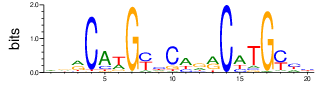

### MA0530.1.png

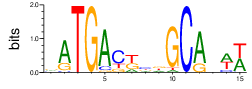

### MA0532.1.png

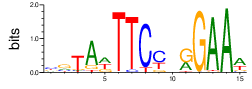

### MA0534.1.png

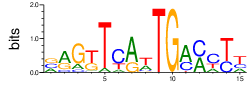

### MA0541.1.png

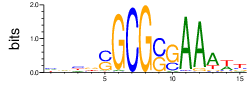

### MA0546.1.png

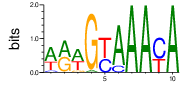

### MA0547.1.png

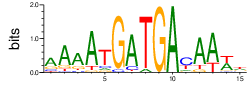

### MA0548.1.png

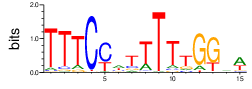

### MA0548.2.png

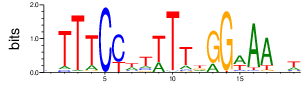

### MA0554.1.png

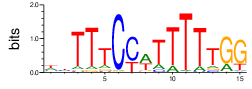

### MA0556.1.png

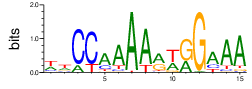

### MA0558.1.png

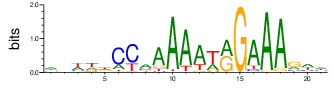

### MA0559.1.png

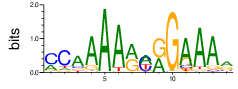

### MA0563.1.png

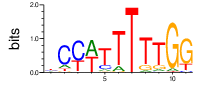

### MA0588.1.png

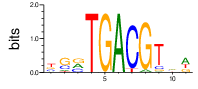

### MA0591.1.png

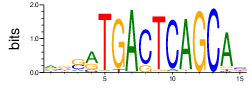

### MA0592.1.png

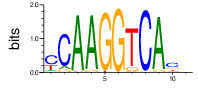

### MA0593.1.png

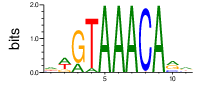

### MA0600.1.png

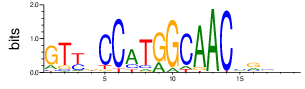

### MA0931.1.png

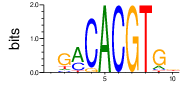

### MA0940.1.png

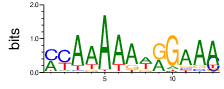

### MA0941.1.png

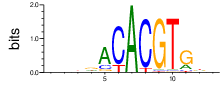

### MA0983.1.png

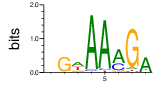

### MA1012.1.png

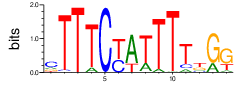

### MA1047.1.png

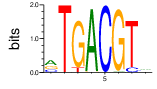

### MA1068.1.png

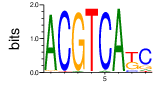

### MA1069.1.png

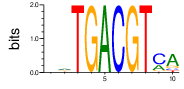

### MA1203.1.png

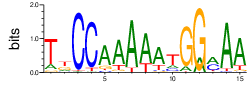

### MA1281.1.png

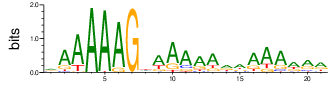

### MA1323.1.png

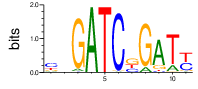

### MA1334.1.png

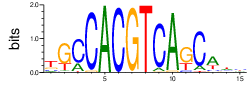

### MA1335.1.png

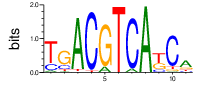

### MA1336.1.png

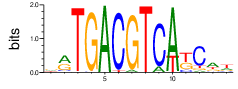

### MA1337.1.png

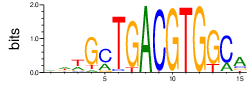

### MA1338.1.png

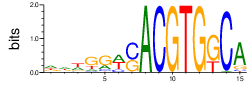

### MA1340.1.png

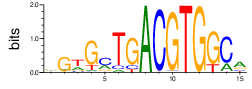

### MA1341.1.png

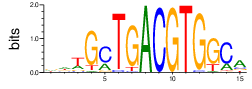

### MA1343.1.png

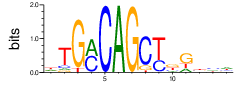

### MA1345.1.png

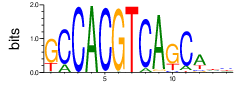

### MA1346.1.png

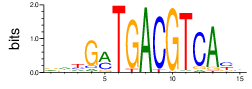

### MA1348.1.png

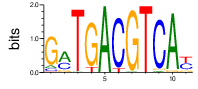

### MA1349.1.png

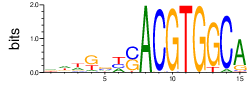

### MA1351.1.png

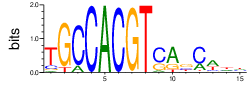
